# Dendritic Cells Overcome Cre/Lox Induced Gene Deficiency by Siphoning Material From Neighboring Cells Using Intracellular Monitoring—a Novel Mechanism of Antigen Acquisition

**DOI:** 10.1101/2023.07.22.550169

**Authors:** Christopher H. Herbst, Aurélie Bouteau, Evelin J. Menykő, Zhen Qin, Qingtai Su, Dunia M. Buelvas, Ervin Gyenge, Neil A. Mabbott, Botond Z. Igyártó

## Abstract

Macrophages and dendritic cells (DCs) in peripheral tissue interact closely with their local microenvironment by scavenging protein and nucleic acids released by neighboring cells. Material transfer between cell types is necessary for pathogen detection and antigen presentation, but thought to be relatively limited in scale. Recent reports, however, demonstrate that the quantity of transferred material can be quite large when DCs are in direct contact with live cells. This observation may be problematic for conditional gene deletion models that assume gene products will remain in the cell they are produced in. Here, we investigate whether conditional gene deletions induced by the widely used Cre/Lox system can be overcome at the protein level in DCs. Of concern, using the human Langerin Cre mouse model, we find that epidermal Langerhans cells and CD11b+CD103+ mesenteric DCs can overcome gene deletion if the deleted gene is expressed by neighboring cells. Surprisingly, we also find that the mechanism of material transfer does not resemble known mechanisms of antigen uptake, is dependent on extra- and intracellular calcium, PI3K, and scavenger receptors, and mediates a majority of material transfer to DCs. We term this novel process *intracellular monitoring,* and find that it is specific to DCs, but occurs in all murine DC subsets tested, as well as in human DCs. Transferred material is successfully presented and cross presented on MHC-II and MHC-I, and occurs between allogeneic donor and acceptors cells—implicating this widespread and unique process in immunosurveillance and organ transplantation.

**ONE SENTENCE SUMMARY:** Dendritic cells maintain RNA and protein levels for conditionally deleted genes by siphoning material from neighboring cells using a novel mechanism.

## INTRODUCTION

Dendritic cells (DCs) are the critical link between innate and adaptive immunity. At steady state, DCs in peripheral tissue scavenge their surroundings for antigen in the form of apoptotic bodies, cell debris, or extracellular vesicles. If they encounter molecules that ligate pattern recognition receptors, they become activated, upregulate costimulatory molecules and MHC-II on their surface, and migrate to lymph nodes to present antigen to adaptive immune cells (*1*, *2*). When cells die or release material into the extracellular environment, DCs are thought to acquire it through multiple endocytic processes, including receptor mediated endocytosis, phagocytosis, or macropinocytosis, which they conduct at high rates (*3*). However, more recent literature has challenged this notion of DCs as scavengers by showing that DCs acquire and cross-present antigen equally well from live cells as they do apoptotic (*4*, *5*). Further, compared to DCs that acquire antigen from apoptotic cells, DCs acquiring antigen from live cells generate larger CD8+ T cell responses and increased protection from lethal tumor challenge when injected *in vivo* (*6*). Separating DCs from live donor cells with a 0.45 μm transmembrane insert prevents cross presentation, indicating live cell contact dependent antigen uptake is critical for inducing an adaptive response (*4*).

Importantly, by acquiring material from live cells, DCs can interact with a variety of molecules that are not usually present in scavenged material. For example, mRNA is degraded as cells go through apoptosis (*7*), and the total volume of material that can be transferred through extracellular vesicles is restricted by their small size. By circumventing these restrictions, DCs can contain large, functionally relevant quantities of RNA and protein from their surroundings. The immunological impact of such transfer has been observed in numerous contexts. Antigen acquired by metallophilic marginal zone macrophages in the spleen is actively transferred DCs, which can promote or suppress adaptive immunity depending on context (*8*). A similar transfer is seen between CXCR1+ macrophages and CD103+ DCs in the context of oral tolerance (9), and macrophages are known to siphon cytosolic material from stem cells during quality control checks (*10*). Our lab found that epidermal Langerhans cells (LCs) can contain KC derived mRNA at nearly 50% the level present in KCs themselves (*11*). Aside from its biological importance, high volume transfer of material from one cell type to another is of concern for researchers utilizing conditional knockout animals.

Cre/Lox animal models are a common tool used to delete genetic regions in specific cell types. This is accomplished by placing expression of the bacterial recombinase Cre under the control of a cell type specific promoter. When expressed, Cre will act on two short LoxP sequences that have been inserted in the gene of interest, resulting in cell type specific gene disruption. However, if DCs can acquire a large enough quantity of material from neighboring cells, the specificity and efficacy of Cre/Lox models may be undermined. Our lab has already shown that DCs can acquire Cre expressed by neighboring cells, potentially resulting in off target effects (*11*), but it remains to be seen whether material transfer can overcome DC specific gene deletion. If so, many DC specific conditional knockout models may be non-functional at the protein level despite successful genetic recombination. Considering the broad usage of such models, further investigation of this concern is warranted.

Of equal intrigue is how DCs are acquiring this material from their neighbors in the first place. Mechanistically, contact dependent material transfer between live cells can occur through trogocytosis, tunneling nanotubes (TNTs), or gap junctions. Trogocytosis, an active “nibbling” process that results in the transfer of surface molecules and membrane fragments (*12*), is routinely used by DCs to acquire peptide:MHC from neighboring cell membranes in a process called cross-dressing, which is important for T cell activation in response to viral infection (*13*). It is also suspected that thymic DCs use trogocytosis to acquire antigen expressed by medullary thymic epithelial cells to help maintain central tolerance (*14*, *15*). TNTs, originally described in 2004 (*16*), are thin, F-actin containing protrusions that enable open ended connection between cells at a distance. Genes required for their formation are highly expressed in many DCs (*17*), and endosomes, viral antigen, and peptide MHC complexes transfer through them during a type one immune response (*18*). Gap junctions enable bidirectional exchange of molecules under 1 kD, including ions, metabolites, small RNAs, and antigenic peptides (19). In some form, all of these mechanisms enable antigen uptake from live cells.

However, considering that trogocytosis predominantly facilitates membrane transfer, *in vivo* evidence of RNA transfer between heterogeneous cell types by TNTs is scarce, and large RNA molecules do not pass through gap junctions, it is difficult to account for the large quantity of material transfer to LCs with them alone. Thus, we also sought to investigate how DCs acquire material from their neighbors. Herein, we show that LCs and CD11b+CD103+ mesenteric lymph node DCs were able to overcome Cre/Lox induced gene deficiency by siphoning RNA and protein from neighboring cells. This ability was shared among all DC subsets tested, exclusive to DCs, and conserved in human DCs. *In vitro*, inhibitors targeting conventional means of antigen uptake failed to prevent siphoning. Instead, DCs used a novel contact-dependent mechanism which we term *intracellular monitoring* (ICM). ICM was dependent on extra and intracellular calcium, and could be blocked with the scavenger receptor inhibitor polyguanylic acid. Material siphoned through this mechanism was presented on MHC-I and MHC-II.

## RESULTS

### DCs can overcome gene deficiency

We previously reported that keratinocytes (KCs) share their gene expression profile with the surrounding Langerhans cells (LCs), affecting LC biology (*11*). Here we tested whether this material transfer could overcome gene deficiencies in LCs. First, we selected two proteins: Cx43 and MHC-II. The rationale behind these proteins was that our previous ATAC-seq data showed that Cx43 genes were open for transcription in KCs, while MHC-II, but not Cx43, was open in epidermal LCs (*11*). Thus, we expected that WT LCs would contain MHC-II, but not Cx43, unless the LCs acquired it from KCs. Flow cytometry analyses of the WT epidermis show that MHC-II is not detected in KCs, but both Cx43 and MHC-II proteins are present in LCs (**Fig. 1A**). The specificity of the anti- Cx43 antibody used here was validated on Cx43 KO cells (clone: CX-1B1; advanced verification by ThermoFisher). Furthermore, ImmGen microarray data corroborated our findings and showed high Cx43 RNA levels in epidermal LCs (**Supplemental Fig. 1A**). Thus, these data confirmed our previous findings that LCs can acquire proteins and RNA from the surrounding cells (*11*). To rule out the possibility that LCs might still express the *Cx43* gene and directly test whether this material transfer can overcome gene deficiency, we bred hLangCre mice to Gja-1^fl/fl^ (Cx43) (*20*) mice, and also used previously characterized hLangCre-MHC-II^f/f^ mice (*21*). The specific genomic recombination of the Cx43 locus in LCs was confirmed using PCR on sorted cells (**Supplemental Fig. 1B**). The flow cytometry staining for MHC-II and Cx43 revealed that while epidermal LCs from Cre+ mice did not have MHC-II, they did have similar levels of Cx43 protein as their Cre-counterparts (**Fig. 1B**), and unchanged levels of Cx43 mRNA (**Supplemental Fig. 1C**). These findings, therefore, support that LCs can overcome a conditional gene deletion if neighboring cells express the missing gene.

**Fig. 1.**
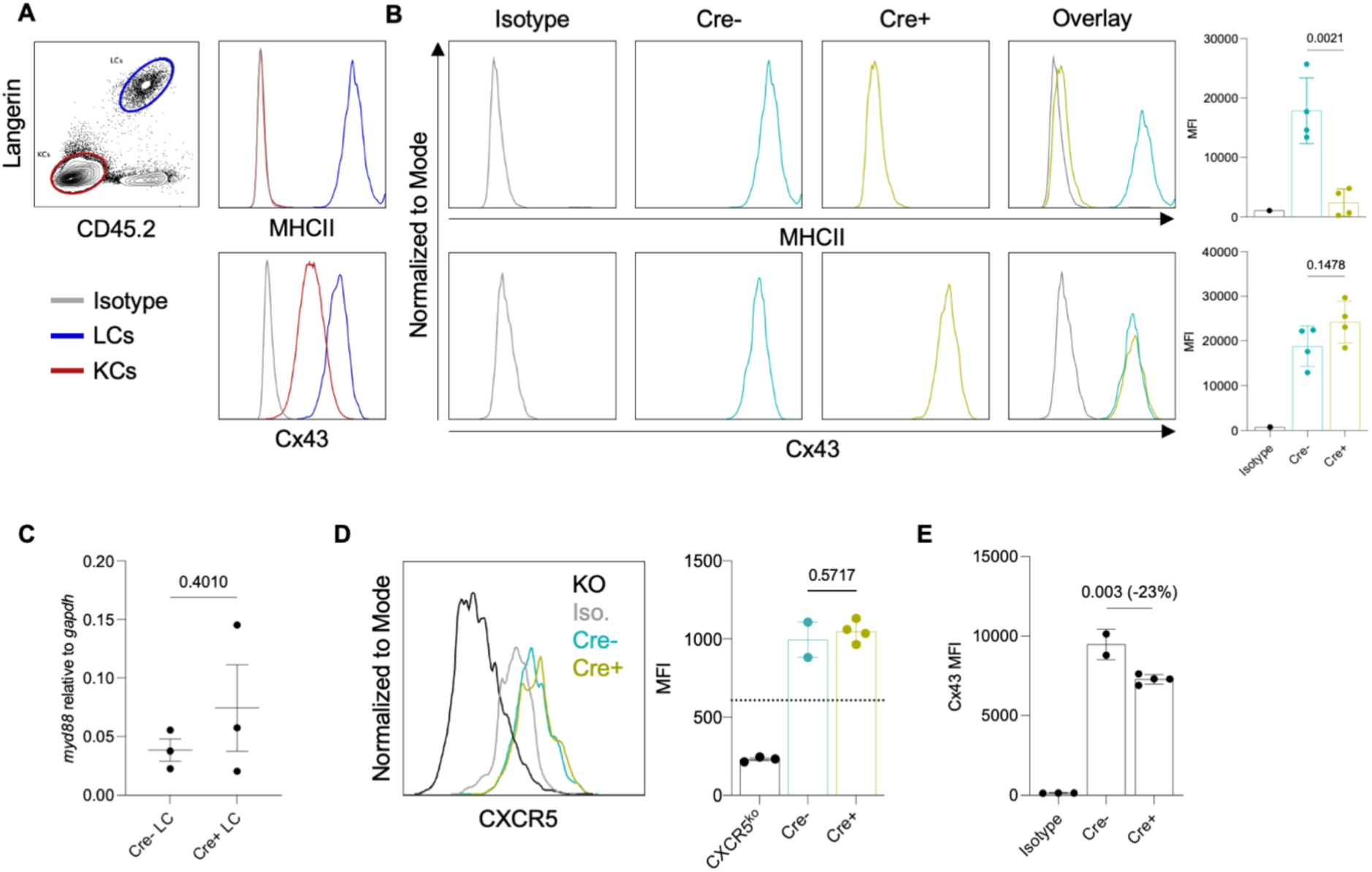
DCs can overcome gene deficiency. (**A**) Gating strategy for identifying keratinocytes and Langerhans cells in an epidermal cell suspension. Epidermal cell suspension from a WT mouse was stained for MHC-II (top) or Cx43 (bottom) or corresponding isotype controls. KCs: keratinocytes, LCs: Langerhans cells. Isotype control signal shown for keratinocytes. Representative flow plots. (**B**) Expression of the indicated proteins by epidermal Langerhans cells derived from MHC-II or Cx43 hLangCre conditional knockout mice. Representative flow plots and summary graphs. Dots represent individual mice. (**C**) *myd88* mRNA level relative to housekeeping gene *gapdh* in either Cre- or Cre+ LCs sorted from hLangCre MyD88^f/f^ epidermis quantified by qPCR. Dots represent individual mice. (**D**) Representative CXCR5 flow staining of splenic DCs from a global CXCR5 knockout mouse (KO), or skin draining lymph node migratory LCs from either Cre- (blue) or Cre+ (green) hLangCre CXCR5^f/f^ mice, or isotype control staining of skin draining lymph node migratory LCs from Cre- hLangCre CXCR5^f/f^ mice, and summary graphs. Dotted line represents isotype control. (**E**) Cx43 flow staining of CD11b+CD103+ mesenteric lymph node DCs derived from either Cre- or Cre+ hLangCre Cx43^f/f^ mice, or isotype control staining of Cre- hLangCre Cx43^f/f^ mesenteric lymph node DCs.

To increase rigor and determine the extent of overcoming gene deficiency by LCs, we further tested the system using two other floxed mouse strains from our hLangCre colony, the hLangCre-MyD88^f/f^ and the hLangCre-CXCR5^f/f^ mice. The hLangCre-MyD88^f/f^ mice were previously published (*22*), while the hLangCre-CXCR5^f/f^ mice were generated for a different project in-house by breeding the hLangCre mice to CXCR5^f/f^ mice (*23*). Both mouse strains showed successful gene recombination in LCs (**Supplemental Fig. 1D and 1E).** Similar to MHC-II and Cx43 protein staining presented above, we sought to assess MyD88 and CXCR5 protein expression levels by flow cytometry. However, the only anti-MyD88 antibody that has been previously reported to generate a positive signal in flow cytometry (MyD88 clone 4D6, Novus Biologicals NBP2-27369) showed WT-level MyD88 staining in global MyD88 KO mice purchased from Jax (**Supplemental Fig. 1F),** rendering it unusable for our purposes. The anti-CXCR5 antibody, which specificity we confirmed on B cells from WT and CXCR5 KO mice (**Supplemental Fig. 1G)**, did not reveal any significant staining in WT epidermal LCs and KCs (**Supplemental Fig. 1G)**. Therefore, to determine whether deletion of MyD88 in LCs can be overcome, we used qPCR on flow-sorted cells. We found that the MyD88 transcripts levels in LCs were not significantly different between the Cre- and Cre+ mice (**Fig. 1C**) and that the KCs had comparable MyD88 to LCs (**Supplemental Fig. 1H**). Thus, these data further confirm that LCs can overcome gene deficiency if localized in a transcript-sufficient niche. Since the epidermal LCs migrate from a CXCR5 deficient epidermal environment to the lymph node, which can be considered as a high CXCR5 niche, we hypothesized that CXCR5 deficient epidermal LCs would become CXCR5+ in the lymph nodes. Indeed, flow cytometry on LCs from hLangCre-CXCR5^f/f^ showed similar levels of CXCR5 as of Cre- littermates (**Fig. 1D**). These findings are in concordance with the previously published observation that LCs in the hLangCre-MHC-II^f/f^ mice acquire MHC-II positivity in the lymph node and induce T cell responses (*24*), and further support our hypothesis that the local environment will dictate whether DCs can overcome gene deficiency.

To determine whether the LCs are unique in overcoming gene deficiencies, we took advantage of the fact that the human langerin promoter in the hLangCre mice drives Cre expression in the CD11b+CD103+ double-positive DCs (**Supplemental Fig. 1I**) in the lamina propria and the mesenteric lymph nodes (*25*). Therefore, next, we assessed the expression of Cx43 in these DCs. As with the skin and skin-draining lymph nodes, we found that double-positive DCs remained CX43 sufficient in the mesenteric lymph nodes (**Fig. 1E**). Thus, these data support that overcoming gene deficiencies is likely universal among DCs.

### Intracellular material acquisition from surrounding cells is specific to DCs and universal among all DC subsets tested

The data above suggest that DCs can acquire cytosolic material from different cell types. To determine which cell types DCs monitor, we used a modified *in vitro* co-culture system that we developed to image RNA acquisition by LCs from epidermal KCs (*11*). Briefly, we co-cultured GFP-expressing MutuDC1 cells (DC cell line with cDC1 phenotypical and functional characteristics) (*26*) with SYTO62-labeled (RNA dye) B-cells, T-cells, peritoneal macrophages, or dermal CD45- cells sorted from adult wild-type naïve C57BL/6 mice (**Supplemental Figure 2A and 2B**). Results show that while DCs acquire RNA from all cell types tested, they most efficiently siphon from macrophages, non-hematopoietic CD45- stromal cells, and keratinocytes (**Fig. 2A**). The RNA transfer was almost entirely unidirectional since the cells were not able to acquire significant amounts of labeled RNA from DCs (**Supplemental Fig. 2C**). Thus, DCs can monitor both hematopoietic and non-hematopoietic cells.

**Fig. 2.**
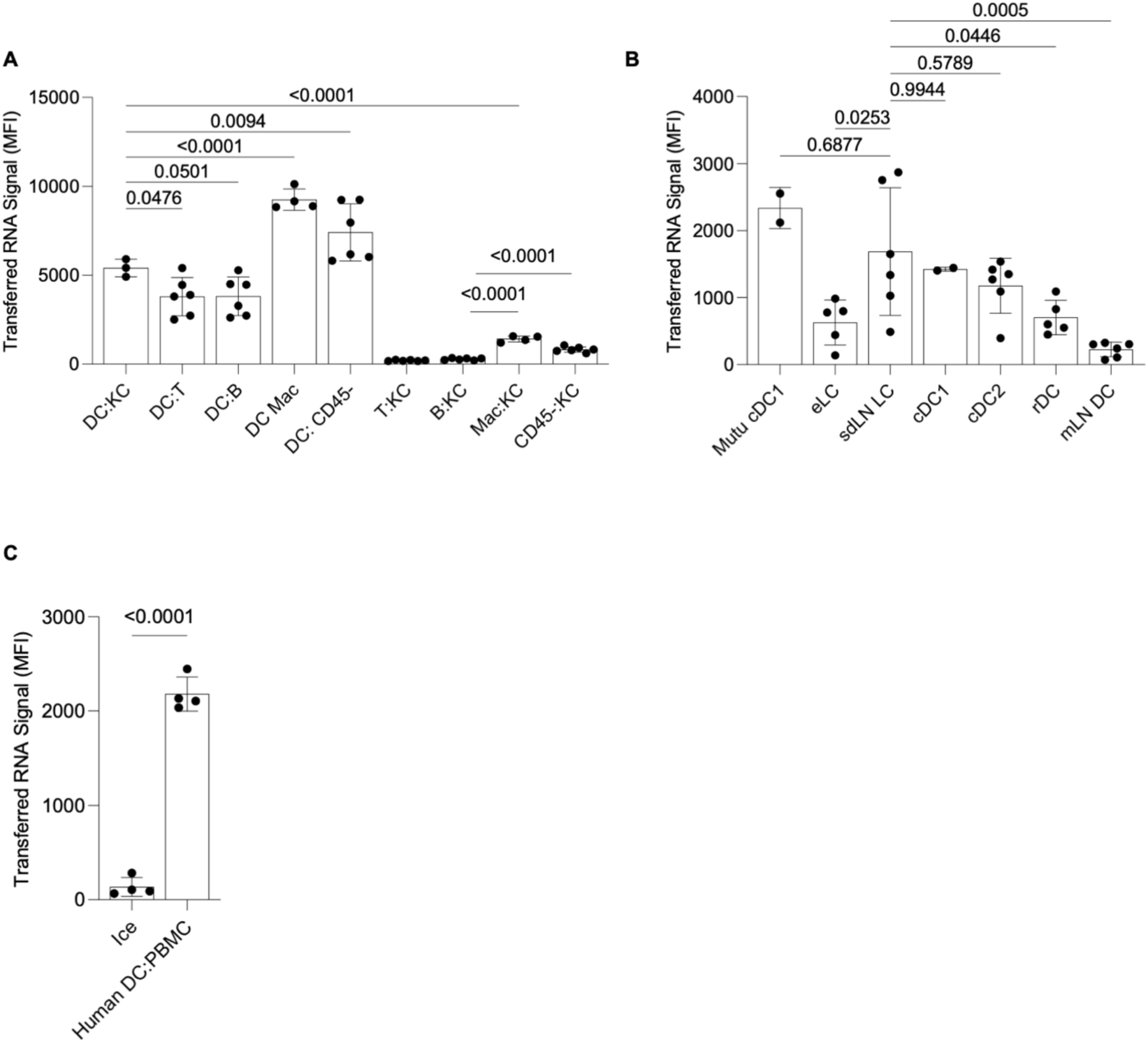
Intracellular material acquisition from surrounding cells is specific to DCs and universal among all DC subsets tested. (**A**) Splenic T cells and B cells, peritoneal macrophages, or dermal CD45- cells were sorted from a wild type C57BL/6 mouse. Cells were labeled with either SYTO62 or CFSE and co-cultured for 45 minutes. Acceptor cells (CFSE labeled) are written first, and donor cells (SYTO62 labeled) are written second (Acceptor:Donor). COCA KCs and MutuDC1s were used as keratinocytes and DCs. Points represent individual replicates derived from two mice. One representative experiment of two shown. (**B**) Epidermal Langerhans cells (eLC), skin draining lymph node migratory Langerhans cells (sdLN LC), sdLN cDC1, sdLN cDC2, resident DCs (rDC), and mesenteric lymph node migratory DCs (mLN DC) were sorted from hLangCre- YFP^f/f^ mice, labeled with CFSE, and co cultured with RNA labeled COCA KCs for 45 minutes. Dots represent individual mice. Data pooled from three experiments. (**C**) Dendritic cells were differentiated from human CD14+ monocytes and labeled with CFSE, then co-cultured with RNA labeled PBMCs on ice or 37°C for 45 minutes. Dots represent individual replicates. One of two repeat experiments is shown.

To better understand the role of intracellular material acquisition, we sought to determine if it is exclusive to DCs, or a property of many cell types. To do that, we co-cultured the sorted T-, B-cells, macrophages and CD45- stromal cells with RNA labeled COCA keratinocytes. We observed that RNA transfer ranged from significantly less efficient (macrophage and stromal cells) to entirely absent (T- and B cells) when using cell types other than DCs as recipients (**Fig. 2A**). This supports that intracellular monitoring is a unique DC property.

We previously reported that DC subsets harbor mRNAs specific to their tissue of residence (*11*). To bring experimental evidence that other DC subsets are capable of intracellular monitoring/surveillance and acquire RNA from the surrounding cells, we sorted DC subsets from single cell suspensions generated from the epidermis, skin draining lymph nodes, and mesenteric lymph nodes of adult naïve mice (**Supplemental Fig. 2D**). DC subsets were then labeled with CFSE and co-cultured with RNA labeled COCA KCs. We found that all DC subsets of the epidermis and skin draining lymph nodes were able to acquire RNA from KCs to varying degrees (**Fig. 2B**), while mesenteric CD11b+CD103+ DCs acquired some, but significantly less RNA. Therefore, these data support that the material acquisition from the target cells by DCs is widespread and not limited to LCs or MutuDC1s.

We previously showed that human LCs, similar to mouse LCs, contain KC-derived keratins (*11*), suggesting that intracellular material acquisition by DCs could be a universal phenomenon. To determine whether human DCs are capable of acquiring RNA from other cells, we differentiated DCs from CD14+ blood monocytes (**Supplemental Fig. 2E**) then incubated them with autologous PBMCs labeled with RNA dye. We found that the DCs were efficient in acquiring RNA from the autologous PBMCs (**Fig. 2C**, **Supplemental Fig. 2F)**. Thus, these data support that intracellular material acquisition by DCs exists in humans and the mechanism is likely preserved.

### Intracellular material transfer is contact dependent, but independent of known antigen acquisition pathways

We previously observed that LCs separated from KCs using 0.4 μm Transwell membrane are unable to acquire detectable levels of KC-derived RNA (*11*). These data suggest that free RNA, exosomes, other forms of extracellular vesicles and/or cell debris that could cross the membrane might not play a significant role in RNA transfer from KCs to LCs. However, some RNA containing microvesicles are larger than 0.4 μm (*27*), and the Transwell membrane may nonspecifically bind some vesicles and therefore hinder their access to LCs. To overcome these caveats and confirm the requirement for physical interaction for RNA transfer from KCs to DCs, we adapted an *in vitro* co-culture system (*28*) where donor cells (KCs) are suspended above recipient cells (DCs) (**Fig. 3A**). In this system, the two cell types are facing each other and are only separated by a thin layer (1.5 mm) of cell culture media. This setup provides DCs unobstructed access to KC-derived exosomes, vesicles, cell debris, and apoptotic cells, while still preventing direct contact. For our system, MutuDC1 cells were seeded at the bottom of a 48-well plate and cells from the murine KC cell line COCA (*29*) were grown on round cover slips, labeled with the nucleic acid dye SYTO62, protein dye cell trace violet (CTV), and suspended using silicone O-rings above the DCs throughout the entire culture period (“physical separation”) (**Fig. 3A and Supplemental Fig. 3A**). Physical separation again prevented transfer of RNA from KCs to DCs as measured by flow cytometry and confocal microscopy, whereas DCs that had direct physical contact with the KCs at 37°C, but not on ice, acquired KC-derived RNA (**Fig. 3B, C, and Supplemental Fig. 3B**). Time lapse imaging shows RNA and protein signal intensity increasing within DCs over time (**Supplemental Fig. 3B**). Imaging further revealed that only DCs in direct contact with donor cells acquired RNA (**Supplemental Video 1**). Together, these data strongly support that released cell vesicles, debris, and apoptotic bodies do not play a significant role in RNA transfer, and they further underpin the need for physical contact for RNA and protein transfer.

**Fig. 3.**
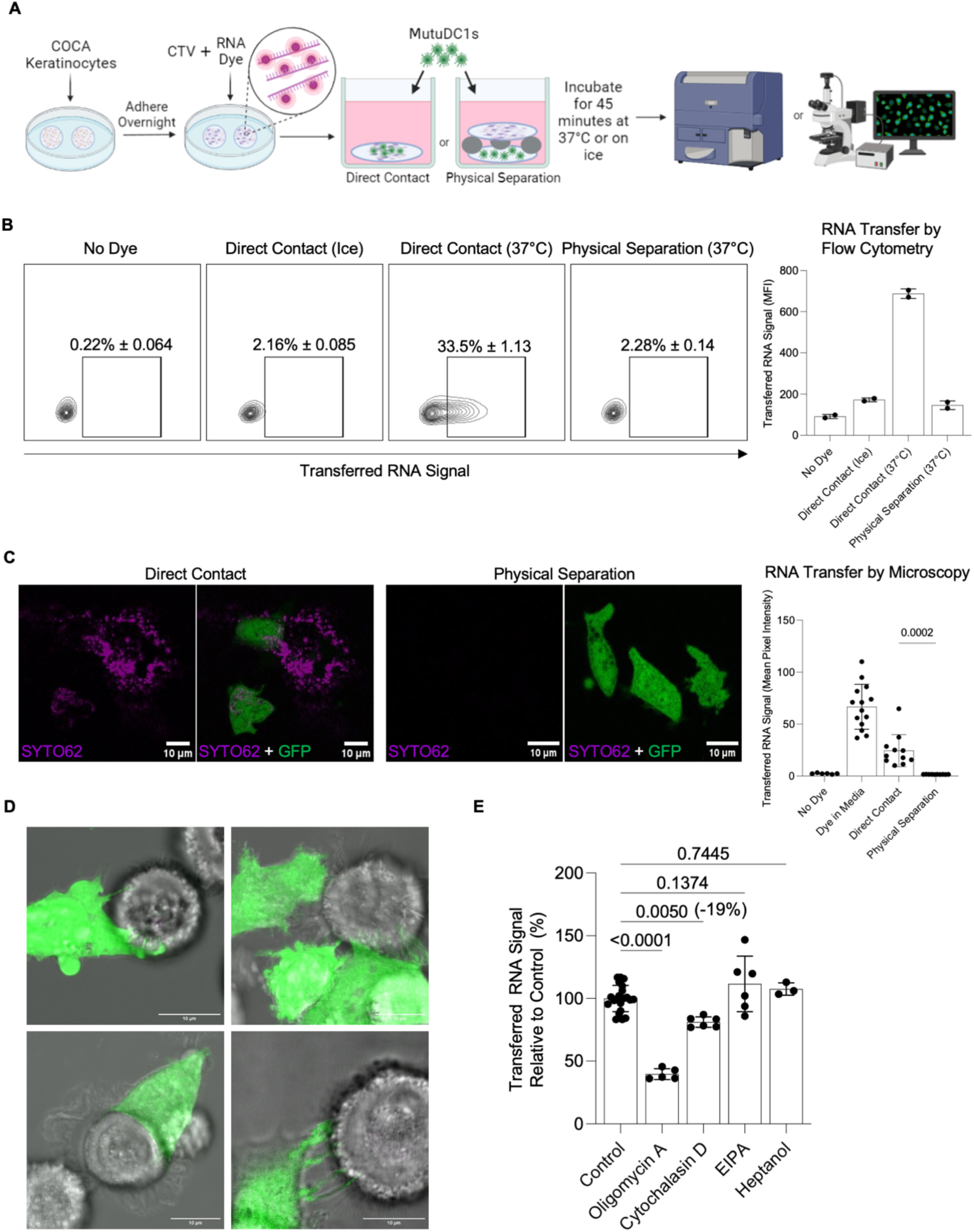
Dendritic cells siphon RNA from neighboring cells through a contact dependent mechanism that does not resemble conventional means of antigen uptake. (**A**) Outline of experiment to measure RNA and protein transfer to MutuDC1s with or without direct contact. (**B**) Flow cytometric analysis of SYTO62 RNA signal measured in MutuDC1s. (**C**) Representative images and ImageJ analysis of SYTO62 signal contained within MutuDC1s in keratinocyte:DC co-cultures. Mean pixel intensity of far-red channel (SYTO62) was calculated within the area occupied by GFP+ MutuDC1s on a per cell basis. Images acquired with a Nikon A1R confocal microscope using a Plan Fluor 40x Oil objective. (**D**) MutuDC1s (green) interacting with keratinocytes. Max projections of Z-stack images taken on a Nikon A1R confocal microscope using a Plan Fluor 40x Oil objective plus 10x scanner zoom. (**E**) Transfer of SYTO62 labeled RNA to MutuDC1s from keratinocytes relative to vehicle controls in the presence of an ATPsynthase inhibitor (1 µM Oligomycin A), an inhibitor of F-actin formation (8 µM Cytochalasin D), a macropinocytosis inhibitor (32 µM 5-(N-Ethyl-N-isopropyl)amiloride (EIPA), and a gap junction inhibitor (5 mM 1-Heptanol). Dots represent individual replicates. Data normalized and pooled from three separate experiments.

To gain insight into the physical interaction and mechanism that allow DCs to acquire cytosolic material from other cells, we co-cultured cells from the murine keratinocyte cell line COCA with MutuDC1 cells and took high resolution confocal images and time-lapse videos of these cells interacting with one another. The physical interaction between the DCs and KCs was diverse in nature, ranging from superficial-looking touching/screening to all the way to DC dendrites pressing into the KC plasma membrane (**Fig. 3D, and Supplemental Videos 2-5)**. Occasionally, DCs formed ring structures when contacting KCs, resulting in RNA containing vesicles. (**Supplemental Fig. 3C, Supplemental Video 3).**

Having established physical interaction as a requirement for material transfer, we next tested whether previously described, standard routes of antigen acquisition, such as phagocytosis, tunneling nanotubes, macropinocytosis, and gap junctions are involved in cytosolic material acquisition by DCs. To do this, we measured RNA transfer by flow cytometry from RNA labeled COCA KCs to MutuDC1 cells during a 45-minute co-culture under different conditions. Transfer was significantly inhibited if they were treated with the ATP synthase inhibitor Oligomycin A (*30*), demonstrating that the process is energy intensive, rather than a passive transfer (**Fig. 3E and Supplemental Fig. 3D**). It was recently established that tunneling nanotubes (TnTs) enable significant RNA transfer between stationary cells (*28*), and we have observed structures resembling TnTs between DCs in some of our long-term (more than 45 minutes) co-cultures (**Supplemental Fig. 3E**). TnTs require intact F-actin (*31*), so we tested for their contribution by using cytochalasin D, an F actin inhibitor (**Supplemental Fig. 3F**). At concentrations of cytochalasin D high enough to inhibit phagocytosis (*32*, *33*), we observed only a minor inhibition (<19%) in RNA transfer (**Fig. 3E**). We next sought to evaluate the contribution of macropinocytosis. 5-(N-ethyl-N-isopropyl)-Amiloride (EIPA), a Na^+^ channel inhibitor known to block macropinocytosis (34). EIPA did not inhibit the RNA acquisition (**Fig. 3E**) but blocked fluorescent dextran uptake by DCs (**Supplemental Fig. 3G**), which is known to be mediated partially by macropinocytosis (*35*). The gap junction inhibitor 1-heptanol (9) also failed to inhibit RNA acquisition (**Fig. 3E**), but did significantly reduce transfer of Calcein dye between COCA KCs (**Supplemental Fig. 3H**), which is partially mediated by gap junctions (*36*). Our findings were not limited to RNA acquisition from KCs. We observed roughly similar responses with an unrelated, cancer cell line, B16 (**Supplemental Fig. 3I**). Therefore, known mechanisms of material uptake such as macropinocytosis, TnTs, phagocytosis, and gap junctions do not appear to play a major role in RNA transfer, and these data are supportive of a novel mechanism.

### DCs acquire cytosolic material from other cells through a novel mechanism dependent on calcium and PolyG blockable receptors

Intercellular interactions are mediated by surface receptors that often rely on Ca^2+^ for binding (*37*). Thus, we next tested whether extracellular Ca^2+^ plays a role in intracellular material acquisition by DCs. We supplemented the DC/KC or DC/B16 co-cultures with 5 mM EDTA to chelate extracellular Ca^2+^. We observed a significant inhibition of material transfer for both DC/KC (**Fig. 4A**, **Supplemental Fig. 4A**) and DC/B16 (**Supplemental Fig. 4A**) co-cultures. The inhibition reached maximum with 5 mM EDTA (**Supplemental Fig. 4B**). To determine whether intracellular Ca^2+^ also plays role in intracellular monitoring, we supplemented the EDTA treated DC/B16 co-cultures or the Ca^2+^-free media with thapsigargin or BAPTA-AM. Adding thapsigargin, a non-competitive irreversible inhibitor of the endoplasmic reticular Ca^2+^ ATPase that is often used to deplete intracellular Ca^2+^ (*38*), or BAPTA-AM, a cell membrane permeable Ca^2+^ chelator, to Ca^2+^- free media had additive effects, leading to an overall 60-70% inhibition of RNA transfer (**Fig. 4B**). Thus, these data suggest a mechanism partially dependent on extracellular and intracellular Ca^2+^. Cadherins and integrins play an essential role in cell adhesion, synapse formation, and intercellular interactions in general, and they are often Ca^2+^ dependent (*37*). Therefore, we next tested the contribution of certain, well-characterized cadherins and integrins to the RNA transfer. The DC/target cell co-cultures were supplemented with blocking antibodies to E-cadherin, CD11b, CD11c, or RGD peptides (to block RGD-binding integrins), or ADH-1 (small molecule inhibitor of N-cadherin). We found no significant inhibition with any of the reagents tested (**Supplemental Fig. 4C**). The binding and potency of the reagents used were confirmed prior to use (**Supplemental Fig. 4D-F, and data not shown**). These data suggest that the integrins and cadherins tested here do not play a substantial role in RNA transfer.

**Fig. 4.**
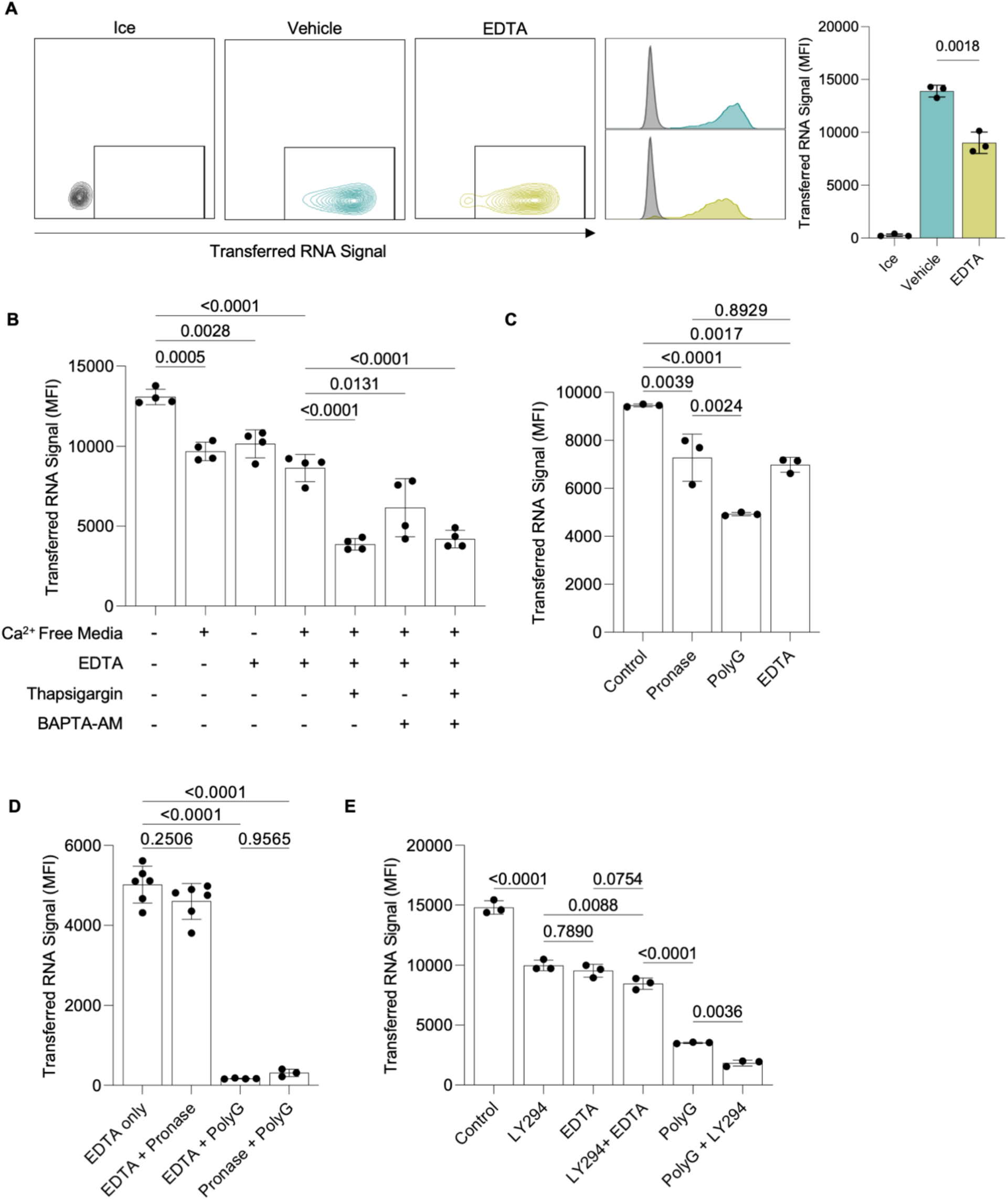
RNA transfer is dependent on calcium and can be partially blocked with the scavenger receptor inhibitor Polyguanylic acid. (**A**) Representative flow cytometry contour plots and histograms of MutuDC1 cells after incubation in direct contact with RNA labeled COCA keratinocytes in the presence or absence of 5 mM EDTA. Dots represent individual replicates. One of three replicate experiments shown. (**B**) RNA dye signal relative to control measured in MutuDC1 cells after incubation with RNA labeled B16 cells. MutuDC1s were treated with thapsigargin (2 µM), BAPTA-AM (50 µM), or both for 30 minutes on ice prior to co-culture in media containing Ca^2+^, Ca^2+^ free media, or 5 mM EDTA. Dots represent individual replicates. One of three repeat experiments is shown. (**C**) RNA dye signal relative to control measured in MutuDC1 cells after incubation with RNA labeled B16 cells. MutuDC1s treated with 32 µg/ml Pronase or co-cultured with B16 cells in the presence or absence of 500 µg/ml PolyG, or 5 mM EDTA as indicated. Dots represent individual replicates. One of three repeat experiments is shown. (**D**) RNA dye signal relative to control measured in MutuDC1 cells after incubation with RNA labeled B16 cells. MutuDC1s treated with 32 µg/ml Pronase or co-cultured with B16 cells in the presence or absence of 500 µg/ml PolyG, or 5 mM EDTA as indicated. (**E**) RNA dye signal relative to control measured in MutuDC1 cells after incubation with RNA labeled B16 cells. MutuDC1s treated for 30 minutes with 50 µM LY294002, 5 mM EDTA, or 500 µg/ml PolyG as indicated. Dots represent individual replicates. One of three repeat experiments is shown.

Protease mixtures, such as Pronase, that can digest a wide range of proteins, are often used to confirm the involvement of cell surface proteins in cellular interactions (*39*). Pronase treatment of DCs has been reported to inhibit cell nibbling/trogocytosis of target cells’ membranes by cleaving off polyguanylic acid (PolyG)-blockable class A scavenger receptor CD204 (*39, 40*). To test whether RNA acquisition by DCs is Pronase sensitive, we treated the MutuDC1s with Pronase as previously described (*39*). The effect of Pronase digestion on cell surface proteins was confirmed by flow cytometry using markers such as CD8 (sensitive), CD11c, CD11b (partially sensitive), and MHC-II (resistant) (**Supplemental Fig. 4D**). Pronase treatment of the DCs caused slight but significant inhibition of the RNA transfer (**Fig. 4C**). In contrast, the inclusion of PolyG with intact DC/B16 co-cultures led to roughly 50% inhibition of RNA transfer (**Fig. 4C**). The difference in percent inhibition indicates that PolyG likely acted through a receptor different than CD204 because this receptor is Pronase sensitive (*39*) and is not expressed by the MutuDC1 cell line used here (**Supplemental Fig. 4G**). Thus, these results suggest that previously described membrane trogocytosis, reliant on Pronase sensitive PolyG blockable scavenger receptor CD204, does not play a significant role in RNA transfer.

Since the Pronase-sensitive receptor(s) showed some redundancy in the material transfer, overlap may exist with the cadherins and integrins tested above that showed no inhibition when tested individually. This possibility prompted us to determine the contribution of those cadherins and integrins to intracellular monitoring in combination with PolyG. Apart from CD11c, we found no additive effects (**Supplemental Fig. 4H**), reinforcing our findings with their single use. PolyG in combination with anti-MHC-II, an abundant surface protein that is not degraded by Pronase (**Supplemental Fig. 4D**), resulted in modest additive inhibition similar to anti-CD11c, indicating this inhibition likely reflects general steric effects and not a specific mechanism (**Supplemental Fig. 4I**) Then we probed whether Ca^2+^ works in concert with the Pronase-sensitive or PolyG blockable receptor. Interestingly, in combination with PolyG but not with Pronase-treated DCs, EDTA almost completely inhibited the RNA transfer (**Fig. 4D**). Pronase treatment in combination with PolyG did however lead to near complete inhibition. PolyG+EDTA inhibition of RNA acquisition was effective with either KCs or B16s as donors (**Supplemental Fig. 4J**), and also inhibited the majority of protein transfer from B16s (**Supplemental Fig. 4K**). PolyG+EDTA also inhibited RNA transfer to human DCs from PBMCs (**Supplemental Figure 4L**). Combining LY294, a PI3K inhibitor, with PolyG, but not EDTA showed additive effect (**Fig. 4D**, **Supplemental Fig. 4M**). Thus, these data suggest a mechanism dependent on Ca^2+^ and that the Pronase-sensitive receptor on DCs is likely Ca^2+^- and PI3K-dependent. Overall, these data support that at least two sets of receptors mediate the RNA and protein acquisition by DCs.

### DCs present the antigen acquired through intracellular monitoring on both MHC-I and MHC-II

Identifying that PolyG with EDTA blocks monitoring with high efficiency allowed us to start addressing the immunological role of this newly discovered antigen acquisition pathway. We first determined whether the acquired antigen is presented on MHC-I. For this purpose, we co-cultured MutuDC1s, known to efficiently cross-present (*26*), or MutuDC2s, unable to cross-present (*41*), with B16 or B16-OVA cells for different time points. Then we determined the presentation of the SIINFEKL peptide by DCs using peptide/MHC-I-specific antibody (**Fig. 5A**). We found detectable levels of SIINFEKL peptides on the MutuDC1s co-cultured with B16-OVA, but not B16, as early as one hour after co-incubation, which increased with time (**Fig. 5B**). In contrast, with MutuDC2s, we failed to detect any significant SIINFEKL presentation at any of the time points tested (**Fig. 5B**). Inclusion of PolyG/EDTA in co-cultures significantly blocked SIINFEKL presentation by MutuDC1s (**Fig. 5C**). The lack of cross-presentation was not due to the failure of MutuDC2 to perform intracellular monitoring, MutuDC2s, albeit less efficient than MutuDC1s, acquired significant amounts of RNA from both B16 and B16-OVA cells (**Fig. 5D**). Thus, these data support that specific DC subsets that are specialized in cross-presentation can process and present the antigen acquired through intracellular monitoring to CD8+ T cells.

**Fig. 5.**
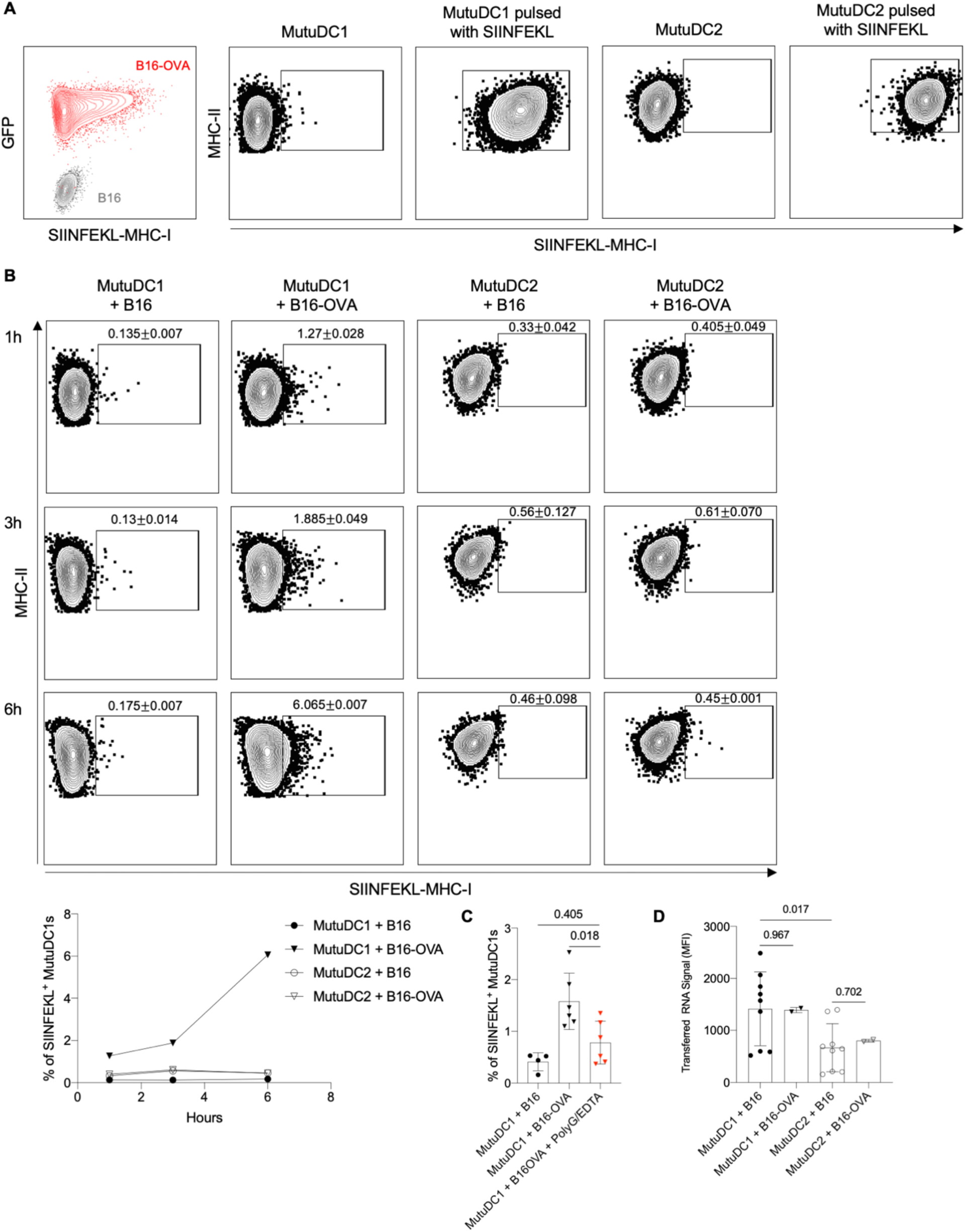
MutuDC1, but not MutuDC2 can cross-present the acquired ovalbumin. (**A**) Control staining with SIINFEKL-MHC-I-specific antibody of B16, B16-OVA, MutuDC1, MutuDC1 pulsed with SIINFEKL, MutuDC2 and MutuDC2 pulsed with SIINFEKL. (**B**) MutuDC1 and MutuDC2 were co-cultured for the indicated time with B16 or B16-OVA and then the SIINFEKL-MHC-I levels determined by flow cytometry. Representative flow plots and summary graph (left lower corner) from one out of two experiments are shown with 2-3 technical replicates. (**C**) As in (**B**), but some of the MutuDC1 co-cultured with B16 or B16-OVA for 3 hours were supplemented with PolyG/EDTA. Data from two independent experiments with 3 technical replicates were pooled. (**D**) MutuDC1 and MutuDC2 were co-cultured with B16 or B16-OVA labeled with SYTO62 for 45 minutes and then the transferred RNA signals (SYTO62) determined by flow cytometry. Data pooled from 3 independent experiments for B16, and one experiment for B16-OVA, with 2-3 technical replicates.

To determine whether the acquired antigens can be presented on MHC-II, we took advantage of the YAe antibody. The YAe antibody recognizes the Ea peptide presented in the context of I-Ab expressed by B6 mice. Ea peptide is derived from BALB/c MHC-II. Thus, we purified T- and B cells from BALB/c skin draining lymph nodes using negative selection, and co-cultured them with B6-derived MutuDC1 and MutuDC2 in the presence or absence of PolyG/EDTA. The rationale behind this setting was that the cells from BALB/c mice would serve as a source of Ea peptide. If the B6 DCs can pick up MHC-II or Ea peptide from the BALB/c cells, process and present it on their MHC-II, then they should turn YAe positive. Indeed, we found that both DC cell lines, although with different efficiency, and dependent on the target cell, became YAe positive (**Fig. 6 and Supplemental Fig. 5**). PolyG/EDTA almost completely blocked the process (**Fig. 6**). Thus, these data support that the antigen sourced through intracellular monitoring can be presented on MHC-II. Furthermore, since this happened in an allogeneic setting, intracellular monitoring might regulate organ rejection.

**Fig. 6.**
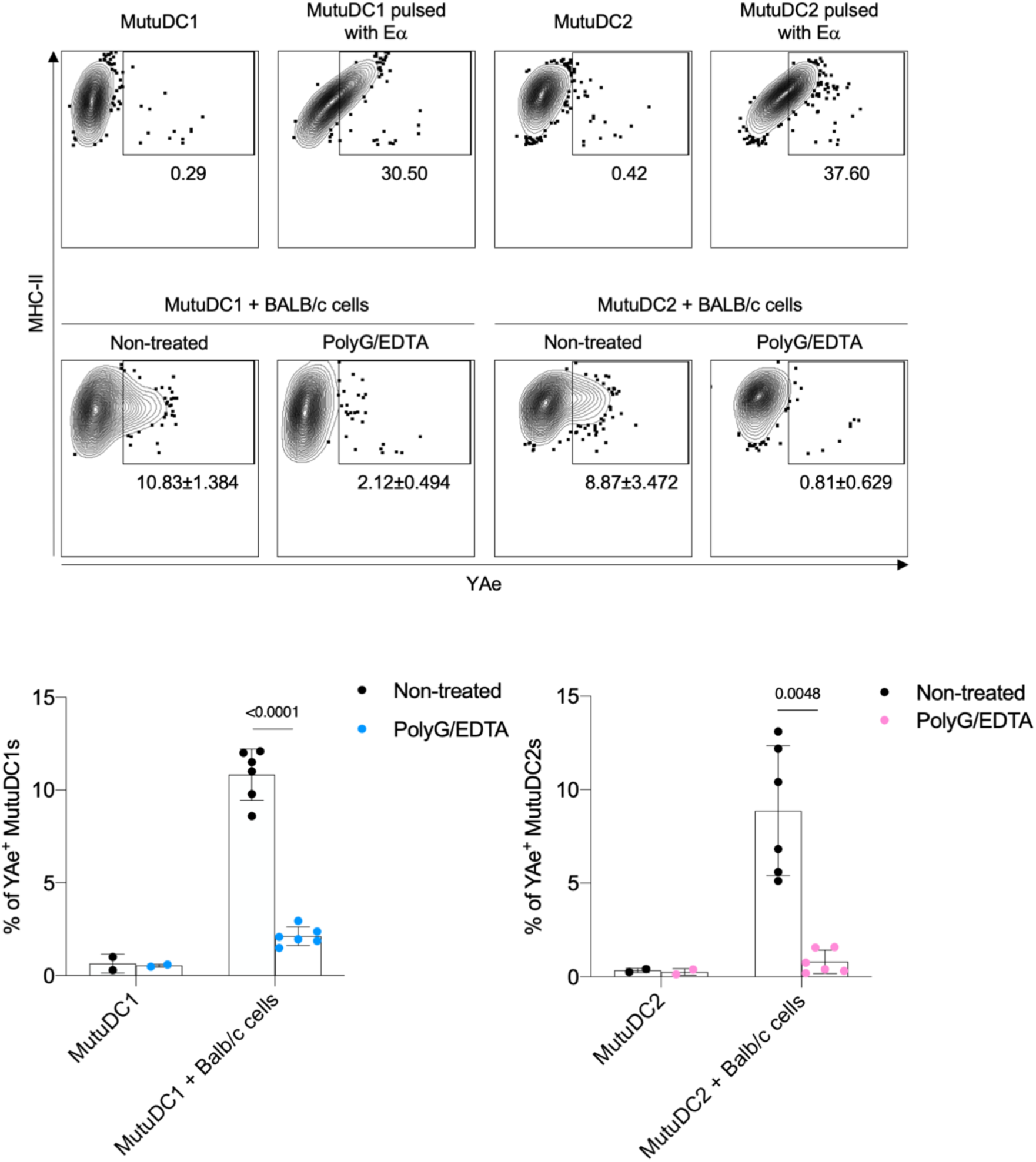
Materials acquired are presented on MHC-II. Top row: Representative flow plots for YAe staining of MutuDC1 and MutuDC2 unmanipulated or pulsed with Ea peptide. Second row: MutuDC1 and MutuDC2 were co-cultured for 3 hours with BALB/c T cells in the absence or presence of PolyG/EDTA and then the Ea presentation of MHC-II determined by flow cytometry. Bottom row: summary graphs for MutuDC1 (left) and MutuDC2 (right). Data from two independent experiments with 3 technical replicates were pooled, except for DC alone controls.

## DISCUSSION

Herein, we showed that epidermal LCs overcome specific gene deletion when neighboring cells contain the missing gene product. Whereas MHC-II gene deletion resulted in a depletion of corresponding protein, gene products of *gja1* and *myd88*— genes that are expressed by neighboring KCs—were present at similar levels in Cre- and Cre+ LCs. CXCR5 and MHC-II deletions are, however, overcome after LC migration to the CXCR5 and MHC-II rich skin draining lymph nodes (*24*). CD11b+CD103+ mLN DCs similarly overcome Cx43 conditional deletion, demonstrating that this trait is likely shared among multiple DC subsets. After *in vitro* co-culture with RNA labeled donor cells, all primary DC subsets tested acquired RNA from neighboring cells to some extent. Co-cultures with different cell types revealed that DCs acquire RNA from a broad range of cell types, but cell types other than DCs either completely fail to acquire RNA, or do so at substantially lower rates. Human DCs were also able to acquire RNA from autologous donor cells. Investigation of the mechanism of RNA transfer revealed it to be dependent on close contact and an active process, as physical separation, or direct contact while on ice results in near complete inhibition of material transfer. Live cell time lapse confocal imaging shows DCs pressing dendrites into the membrane of donor cells and maintaining close contact. Actin cytoskeletal inhibition, which is known to prevent most forms of phagocytosis, endocytosis, trogocytosis, and the formation of tunneling nanotubes (*33*, *42–46*), does not substantially prevent transfer. RGD peptide or blocking antibodies against integrins commonly involved in endocytic process such as CD11c and CD11b (*43*) also have no effect on transfer. EIPA, an inhibitor of macropinocytosis, and 1-heptanol, a gap junction inhibitor, also fail to prevent transfer. Instead, we find that transfer is partially inhibited by removing calcium from the media, PI3K inhibition, or by the introduction of PolyG into co-cultures. Combining PolyG with EDTA or Pronase treated DCs, but not EDTA with Pronase treated DCs, is sufficient to block most of the transfer. Transferred material is successfully presented and cross presented on MHC-II and MHC-I, and occurs between allogenic donor and acceptor cells. Due to its discordance with conventional means of antigen uptake, we termed this mechanism as *intracellular monitoring* (ICM).

The observation of inaccessible *gja1* chromatin in epidermal LCs (ATAC-seq) (11) but high *gja1* transcript levels strongly suggests that LCs acquire RNA for this gene from another source. Here, we show this definitively by specifically deleting *gja1* in LCs and observing no deficit in RNA or protein in knockout cells. Considering that a deficit is observed after specific deletion of the LC specific protein MHC-II, the most likely explanation for how LCs acquire missing RNA and protein is that they take it from neighboring cells. Cre expression in mLN DCs and the use of multiple knockout models show this remains the case for multiple DC subsets and proteins. Critically, this greatly raises the probability of type II error in studies utilizing conditional knockout models targeting DCs, as the acquisition of protein from neighboring cells may lead researchers to incorrectly conclude their knockout has no effect. Furthermore, these data raise serious concerns regarding gene expression databases on DCs, which based on our data, are likely a snapshot in time and a mixture of self and acquired mRNA from the local cells. Thus, these findings highlight the importance of verifying target protein depletion, not just successful genetic recombination, and the need for curation of RNA-seq data.

Aside from the immediate practical concerns surrounding ICM and conditional knockouts, ICM may also be relevant to important biological functions such as microenvironmental adaptation, immunosurveillance, and tolerance. Immune cell adaptation to the local microenvironment is a concept that has been extensively studied in macrophages, and refers to the dramatic shift in the chromatin landscape in macrophages in response to environmental queues such as retinoic acid or heme (*47*). These changes endow macrophages with functions necessary to operate properly in their local niche, and contribute to, instead of interrupt, the function of their resident organ (*48*). ICM may be providing a similar benefit to epidermal Langerhans cells. While we did not directly test the functionality of transferred protein in this study, one out of many viable explanations for LC’s possession of Cx43 could be to prevent the disruption of wound healing, which is dependent on the direct transfer of Ca^2+^, IP_3_, and ATP through gap junctions and the ensuing calcium waves (*49, 50*). These waves are projected to travel through LCs as well as KCs (*51*), supporting that LCs might acquire functional protein to help them adapt to their environment and possibly to be alerted by infected KCs. Further studies investigating the fate and function of transferred material will help elucidate the roles of ICM in microenvironmental adaptation.

Considering DCs overcome deficiency of multiple proteins, including ones they express on their own, it is likely that DCs continually and non-specifically conduct ICM. Among cell types tested, ICM was specific for DCs, and, to a lesser degree, macrophages. Its specificity for antigen presenting cells, combined with the finding that acquired protein is presented and cross presented on MHC-I and MHC-II, points toward ICM being highly relevant to typical DC functions such as immunosurveillance and tolerance, and may explain the long-standing mystery of how DCs receive material from other cells for cross-presentation (*4*, *5*, *52*). DCs can monitor all donor cells tested, but more efficiently monitor CD45- cells and macrophages. If DCs use ICM to detect pathogens, monitoring macrophages with high efficiency would provide an evolutionary advantage, as these cells are often the first to encounter pathogens, and are more likely to contain a diverse pool of antigen. The low monitoring efficiency of the CD103+ cDC2 (CD11b+CD103+) mesenteric DCs, which migrate from the predominantly tolerogenic environment of the gut and are involved in T_reg_ and T_h17_ cell induction (*2*, *53*), supports this and argues against ICM playing a critical role in maintaining tolerance, though this cannot be ruled out. Of note, low mLN DC monitoring efficiency aligns well with their lesser ability to overcome Cx43 knockout relative to epidermal LCs. On the other hand, it is also possible that while they are in the lamina propria of the gut, the very same DCs might possess high ICM capability, then downregulate it by the time they reach the mesenteric LNs to protect the cargo that requires tolerance induction. While this remains to be experimentally tested, we found that the opposite is true for LCs. LCs that have migrated to LNs are more efficient in ICM than their peripheral counterparts in the epidermis. Whether these site-specific differences have evolved to better serve tolerance induction or simply reflect that ICM is a tool for pathogen detection or that the monitored cell type and environment in the periphery will imprint a downstream program in the DCs, remains to be addressed.

From an evolutionary standpoint, it is logical that DCs would use ICM to detect pathogens. Intracellular pathogens have evolved complex and effective mechanisms to interfere with host cell processes and limit detection by the host immune system (*54–56*). Relying on material released or presented by infected cells is therefore not a dependable way to detect meddling pathogens. Direct presentation after infection also cannot be relied on, as not all viruses are DC tropic or highly cytopathic, and, even if they are, DCs themselves could be subjected to pathogen immune evasion mechanisms, resulting in inefficient presentation. Thus, ICM may potentially counteract some of the evading mechanisms developed by the pathogens. Further, if ICM facilitates inflammatory immune responses, our observation that DCs perform it in allogeneic and xenogeneic (unpublished observation) settings suggests it may play a role in organ rejection and be a valid therapeutic target.

While the exact mechanism of ICM remains to be determined, experiments conducted herein sufficiently differentiate it as a novel process. Its contact dependent nature rules out the uptake of extracellular material as a major contributing factor of RNA and protein transfer. Phagocytosis of dead or dying cells can be ruled out as dying cells are not prevented from detaching and coming in contact with DCs in our physical separation experiments. Cultured cells also maintained high viability throughout experimentation, and no donor cells or debris were observed in contact with DCs in physical separation experiments. TnTs are notoriously fragile and can be eliminated with doses as low as 50 nM Cytochalasin D (*57*), ruling out their involvement in material transfer. While some aspects of intracellular monitoring are reminiscent of trogocytosis, our findings are not consistent with this mechanism. Trogocytosis has been successfully inhibited by PolyG in DCs (*39*) and LY294002 in other cell types (*43*), similar to what we observed, however Harshyne et al. show that PolyG inhibits trogocytosis through blockade of Scavenger receptor A (CD204) (*39*), which the MutuDC1 cells used in this study do not express. Further, the actin cytoskeleton is required for trogocytosis, whereas cytochalasin D fails to substantially prevent intracellular monitoring. Finally, trogocytosis almost exclusively refers to the transfer of membrane between cells (*58*), not the transfer of cytosolic material, further differentiating our observation of RNA and protein transfer from trogocytosis. Considering their size limitations, it is very unlikely that gap junctions would enable substantial RNA and protein transfer. Interestingly, in their study of oral tolerance, Mazzini et al. find that gap junctions only partially mediate transfer from macrophages to DCs, and note that “still-unknown mechanisms” may be contributing (*9*). In retrospect, it is likely that at least some of the material transfer from macrophages to DCs reported by Mazzini et al., was through ICM.

We previously reported that human LCs, like their mouse counterpart, also contain detectable levels of Krt14 mRNA (*11*). Here, we further showed that human DCs differentiated from CD14+ monocytes efficiently acquire RNA from PBMCs, and that RNA transfer can be significantly inhibited by PolyG/EDTA. These data support the translatability of our mouse data and indicate ICM may be a conserved process.

In summary, we show that a widely used research tool—Cre/Lox conditional gene knockout—may be inherently flawed when applied to DCs due to a novel antigen uptake process that we term intracellular monitoring.

## MATERIALS AND METHODS

### Study design

This study aimed to determine how DCs acquire cytosolic material from surrounding cells and define roles for this process. We designed and performed experiments using cellular immunology techniques, flow cytometry, qPCR, immunofluorescence microscopy, murine *in vivo* and *in vitro* models, and human models. The sample size and number of independent experiments are indicated in each figure legend.

### Ethics statement

Institutional Care and Use Committee at Thomas Jefferson University approved all mouse protocols. Protocol number: 02315.

### Mice

hLangCre-YFP^f/f^ (*59*), hLangCre-MHC-II^f/f^ (*21*) and hLangCre-MyD88^f/f^ (*22*) mice were previously described. hLangCre-Gja1^f/f^ (Cx43) and hLangCre-CXCR5^f/f^ mice were generated in house by crossing the huLangCre mice with Gja1^f/f^ mice (JAX stock#008039) (*20*) and CXCR5^f/f^ (*23*), respectively. All experiments were performed with 8–12-week-old female and male mice. Mice were housed in microisolator cages and fed autoclaved food.

### Confirmation of genetic recombination

Epidermal cell suspensions were generated from hLangCre-Gja1^f/f^, hLangCre-MyD88^f/f^, and hLangCre-CXCR5^f/f^ mice as previously described (*60*) and stained with eBioscience’s Fixable Viability Dye eFluor780, CD207 (4C7), I-A/I-E (M5/114.15.2), CD45.2 (104). LCs were sorted as live CD207+MHC-II+CD45.2+ cells, and KCs as triple negative. DNA was extracted and genetic recombination verified using primers CTTTGACTCTGATTACAGAGCTTAA (forward) and GTCTCACTGTTACTTAACAGCTTGA (reverse) for *gja1*, which amplify a 600 bp segment in non-recombined Cre- mice, and no band in recombined mice. *myd88* genetic recombination was verified using primers GGGAATAATGGCAGTCCTCTCCCAG (forward) and CAGTCTCATCTTCCCCTCTGCC (reverse) for *myd88*, which amplify a 400 base pair segment in recombined cells. *cxcr5* genetic recombination was verified using primers AGGAGGCCATTTCCTCAGTT (forward), GGCTTAGGGATTGCAGTCAG (reverse), and TTCCTTAGAGCCTGGAAAAGG (recombination), which amplify a 292 base pair segment in recombined cells or a 375 base pair product in non-recombined cells.

### Cell lines and culture conditions

The COCA, murine epidermal keratinocyte cell line derived from the back skin of adult C57BL/6 mice (*29*) was purchased from ECACC (Salisbury, UK, #10112001). Cells were cultured in CnT-07 media (Fischer Scientific, #NC9474150) in polystyrene tissue culture treated flasks. At 80-90% confluence, cells were removed from the flask by incubation with Trypsin-EDTA (0.25%) (Fisher Scientific, #25-200-072) for 8 minutes at 37 °C. Trypsin was then diluted with a 1:1 dilution of seeding media (EMEM, Caisson Labs), 4% FBS pre-treated with Chelex resin (Bio-Rad, #1421253; 0.2 mM CaCl_2_). Cells were centrifuged at 240 g and split at 1:3 into a new flask containing seeding media. Cells were left in seeding media for 6 hours to promote attachment. Seeding media was then removed, cells were washed twice with PBS, and media was replaced with CnT-07. Cells were not used for experiments before reaching total confluence.

The MutuDC1 and MutuDC2 DC cell lines, isolated from a C57BL/6 murine tumor, were a generous gift from Prof. Hans Acha-Orbea (26, 41). The cell lines were cultured in polystyrene tissue culture treated flasks in IMDM, supplemented with glutamax, 8% FBS (Fisher Scientific, #MT35010CV, Lot: 14020001), Pen-Strep, and 55 µM 2-ME, and maintained in a humidified incubator at 37 °C and 5% CO_2_. To remove the cells from the flask, the media was aspirated and replaced with 5 mM EDTA and incubated at room temperature for 10 minutes. Media was then added to the flask at a 1:1 ratio and cells were centrifuged at 390 g. The cells were then resuspended in media and split at 1:5.

The B16-F10 murine melanoma cell lines were a generous gift from Dr. Michael Gerner, University of Washington. Cells were cultured in polystyrene tissue culture treated flasks in RPMI supplemented with 10% NBCS (Fisher Scientific, #26010074), Na-Pyruvate, NEAA, L-Glutamine, Pen-Strep, and 55 μM 2-ME, and incubated at 37 °C and 5% CO_2_. At ∼90% confluence, cells were harvested by incubation with Trypsin-EDTA (0.25%) for 3 minutes and split at 1:10.

### MutuDC1/COCA keratinocyte coverslip co-culture experiments

COCA keratinocytes were allowed to adhere to 8 mm coverslips (Electron Microscopy Sciences, #72296-08) overnight in seeding media. Seeding media was then removed and cells were washed twice with PBS before media was replaced with CnT-07 and incubated overnight again until cells were nearly confluent. The following day, CnT-07 media was replaced with HBSS containing 50 nM SYTO62 RNA dye (Fisher Scientific, #S11344), and cells were incubated for 20 minutes at 37 °C before being washed twice with HBSS. Coverslips with stained keratinocytes were carefully moved using forceps to the bottom of a 48 well plate with cells facing up, and 25,000 MutuDC1 cells were added directly on top. Alternatively, an autoclaved silicone O-ring (width 1.5 mm) was placed in the well, and 25,000 MutuDC1s were added inside of it and allowed to settle. Keratinocyte coated coverslips were then placed on top of the O-ring with cells facing down. Wells were filled with DC media so that suspended coverslips were completely submerged. After a 45 minutes incubation at either 37 °C or on ice, MutuDC1 cells were resuspended by pipetting up and down, and transferred RNA was measured by flow cytometry. The same protocol was conducted using ibidi 8 chamber slides (Fisher Scientific, #NC0704855) to allow for confocal imaging. Images were taken on a Nikon A1R Confocal microscope using a Plan Fluor 40x Oil objective at the end of the 45 minute incubation. The amount of transferred RNA was measured in ImageJ by calculating the mean far-red pixel intensity (SYTO62 signal) contained within regions of high green channel signal (representative GFP^+^ MutuDC1s). Briefly, green channel images were converted to 8-bit and thresholded appropriately. Watersheding was used to parse clumped cells, then the analyze particle’s function was used to identify Regions of Interests corresponding to area within MutuDC1 cells. These ROIs were then applied to the far-red channel and mean pixel intensity was calculated.

### Study of RNA transfer in the presence of inhibitors

Once nearing confluence, MutuDC1, B16s, or COCA cells were harvested as described and suspended in 1 ml of their respective media. Donor cells – either COCA cells or B16s -- were pelleted and resuspended at 10^6^ cells/ml in pre-warmed HBSS containing 200 nM SYTO62 dye, then incubated at 37 °C for 20 minutes. Donor cells were then washed twice with ice cold PBS and resuspended in DC media with inhibitor or vehicle. At this time, MutuDC1 cells were also resuspended in media containing inhibitor or vehicle. Cells were protected from light, then left on ice for 30 minutes in the presence of inhibitor. Reagents used in this study include: 8 µM Cytochalasin D (Millipore Sigma, #250255), 10 µM Oligomycin A (Selleck Chemicals, #S1478), 32 µM 5-(N-Ethyl-N-isopropyl)amiloride (EIPA) (Sigma Aldrich, #A3085), 5 mM 1-Heptanol (Fisher Scientific, #AAA12793AE), 5 mM EDTA (Fisher Scientific, #AM9260G), 2 µM Thapsigargin (Fisher Scientific, #NC9006970), 50 µM BAPTA-AM (Fisher Scientific, #50-201-0390), 50 µM LY294002 (Tocris, #1130), 350 nM ADH-1 (MedChem Express, #HY-13541), 500 µg/ml Polyguanylic acid (Sigma-Aldrich, #P4404), and 1 mg/ml RGD peptide (Selleck Chemicals, #S8008). Blocking antibodies specific for the following proteins were used: 2.5 μg/ml CD11b (M1/70, BioLegend, #101224), 2.5 μg/ml CD11c (N418, BioLegend, #117322), 1 μg/ml CD204 (M204PA, Fisher Scientific, #12-2046-80). In some cases, MutuDC1 cells were treated with 32 μg/ml Pronase (EMD Millipore, #53702) for 20 minutes at 37 °C as previously described (*39*). Both COCA and MutuDC1 cells were then counted and combined into a 96-well plate containing media with inhibitor or appropriate vehicle. Eighty thousand COCA keratinocytes or B16 cells were combined with 10,000 MutuDC1s. In some experiments, primary cells were used for co-culture after being sorted as described. In these experiments, 10,000 acceptor cells were stained with 5 µM CFSE (Fisher Scientific, #50591407) for 5 minutes on ice and mixed with 80,000 donor cells. Once plated, cells were mixed and moved directly from ice to a 37 °C, 5% CO_2_ incubator, and incubated for 45 minutes. After incubation, cells were moved back to ice, resuspended by pipetting, filtered through a 50 µm filter, and run on flow cytometer.

### Live cell imaging using confocal microscopy

Cultures were imaged in ibidi 8 chamber slides held in a humidified chamber at 37 °C and 5% CO_2_.

### Cross-presentation experiment

B16 or B16-OVA cells were co-cultured with MutuDC1s or MutuDC2s as triplicates in a U-bottom 96-well plate in a CO_2_ incubator for 1, 3 and 6 hours. Some of the 3 hours cultures were supplemented with a standard dose of PolyG/EDTA. After incubation the cells were washed, stained with eBioscience’s Fixable Viability Dye eFluor780, I-A/I-E (M5/114.15.2) and SIINFEKL/MHC-I (eBio25-D1.16) and analyzed by flow cytometer.

### Presentation on MHC-II

BALB/c T and B cell purified by negative selection following the instruction provided by BioLegend and Stem Cells, respectively, were co-cultured with MutuDC1s or MutuDC2s for 3 hours in the absence or presence of standard dose of PolyG/EDTA. After incubation the cells were washed, stained with eBioscience’s Fixable Viability Dye eFluor780, I-A/I-E (M5/114.15.2) and YAe (eBioY-Ae) antibodies and analyzed by flow cytometer.

### Cell sorting

T and B cells were sorted from the spleen of wild-type C57BL/6 mice. A single cells suspension was generated and stained with eBioscience’s Fixable Viability Dye eFluor780 and anti-CD90.2 (30-H12, BioLegend, #105308) and anti-CD19 (6D5, BioLegend, #115546). CD90.2+ cells were considered T cells and CD19+ cells were considered B cells. Cells were sorted on a BD FACSAria II sorter.

CD45- cells were sorted from the dermis of a wild-type C57BL/6 mouse. A single cell suspension was generated and stained with eBioscience’s Fixable Viability Dye eFluor780 and anti-CD45.2 (104, BioLegend, #109808).

Lymph node cell suspensions were generated from hLangCre-YFP^f/f^ mice and enriched for CD11c+ cells using EasySep™ Mouse CD11c Positive Selection Kit (StemCell, #18758). Post enrichment cells were stained with eBioscience’s Fixable Viability Dye eFluor780 and the following antibodies: CD11c (N418), I-A/I-E (M5/114.15.2), CD103 (2E7) and CD207 (4C7). Resident DCs were gated as live CD11c^hi^ MHCII^med^, migratory LCs were gated as live MHCII^hi^CD11c^med^YFP+, cDC1 were gated as live MHCII^hi^CD11c^med^CD103+CD207+, cDC2s were gated as live MHCII^hi^CD11c^med^CD207-, mesenteric DCs were gated as live YFP+. Cells were sorted on a BD FACSAria II sorter.

Epidermal cell suspensions were generated from hLangCre-YFP^f/f^ mice. Cell suspensions were stained with anti-CD45.2 (104) PE antibody and enriched for LCs using EasySep™ PE Positive Selection Kit II (StemCell). Post enrichment cells were stained with eBioscience’s Fixable Viability Dye eFluor780 and sorted for live YFP+ cells on a BD FACSAria II sorter.

Cells from a peritoneal lavage were stained with viability eBioscience’s Fixable Viability Dye eFluor780, F4/80 (BM8), and CD11b (M1/70). Macrophages were gated as F4/80^hi^CD11b^hi^ cells and sorted on a BD FACSAria II sorter.

### Quantitative PCR

Cells were sorted directly into 100 μl of lysis buffer supplemented with beta-mercaptoethanol from the Agilent Absolutely Rna Nanoprep Kit (Neta Scientific, #400753) and RNA was extracted according to the manufacturer’s protocol. Iscript Reverse Transcription supermix (Bio-Rad, #1708841) was used to generate cDNA. qPCR reactions were conducted using 1 μl of cDNA product in 10 μl total reaction volume using Itaq Universal SYBR green (Bio-Rad, #1725121) with primers at a concentration of 500nM. Primers used are as follows: IDT *myd88* Primetime primers (IDT, Mm.PT.58.33389595), *gapdh* Reverse: TCTTGCTCAGTGTCCTTG, *gapdh* Forward: CTTTGTCAAGCTCATTTCCTGG, *gja1* reverse: CGTGGAGTAGGCTTGGAC *gja1* forward: TTCCTTTGACTTCAGCCTCC.

### Generation of human DCs and co-culture with autologous PBMCs

To make human myeloid-derived DCs, 2×10^6^ human CD14+CD16- blood monocytes isolated with EasySep™ Human Monocyte Isolation Kit (StemCell) were cultured in six-well plates (2 ml per well) in complete RPMI 1640 medium + 10% FBS + 10 ng/ml human IL-4 (PeproTech) + 100 ng/ml human GM-CSF (PeproTech). Half of the medium was changed at day 2 and at day 4, maintaining the same concentration of IL-4 and GM-CSF. Immature DCs were collected on day 5 and analyzed using a flow cytometer (CD14, HLA-DR and CD86). After confirmation the remaining DCs were used for intracellular monitoring. Briefly, the DCs were labeled with CFSE and co-cultured with autologous PBMCs labeled with RNA dye in the presence or absence of PolyG/EDTA for 45 minutes. The RNA acquisition by DCs from PBMCs was confirmed by flow cytometer.

### Actin polymerization inhibition assay

100,000 MutuDC1 cells were plated in each chamber of a removable 8 chamber slide and left to incubate for 2 hours at 37 °C to allow attachment. Media was then aspirated off and replaced with media containing 4 μM Cytochalasin D or DMSO control and incubated for an additional 30 minutes. The media was then aspirated off, and cells were washed once with PBS and fixed with 4% paraformaldehyde (EM Grade, Fisher Scientific, #50-980-495) in PBS for 20 minutes. After fixation, the paraformaldehyde was removed, and cells were washed two more times with PBS. 183 nM Phalloidin labeled with rhodamine working solution was then added, and cells were incubated for 60 minutes at room temperature. The cells were washed twice more with PBS. The removable chambers were detached, and coverslips placed over the cells using mounting media for imaging.

### TRITC-Dextran uptake

MutuDC1 cells were incubated in 32 μM EIPA or vehicle control for 30 minutes on ice. Cells were then centrifuged and resuspended in HBSS containing 16 μM or 32 μM EIPA or vehicle and 20 μg/ml 10 KDa TRITC-Dextran (Fisher Scientific, #D1868) and incubated at 37°C for 30 minutes. After incubation, cells were washed twice with PBS, then resuspended in Trypsin-EDTA (0.25%) for 5 minutes to remove any Dextran adherent to the outside of the cell. Cells were then washed twice more and analyzed by flow cytometry.

### Gap junction Calcein transfer assay

COCA keratinocytes were separated into two aliquots. One aliquot was stained with 10 μM Tag-it Violet (BioLegend, #425101) diluted in HBSS for 5 minutes on ice at a cell concentration of 1 million cells per milliliter. After incubation, 5x volume of serum containing media was added to the tube and cells were incubated for an additional 5 minutes on ice. Cells were then centrifuged at 240 g for 7 minutes, washed once with PBS, and finally resuspended in media. The second aliquot of cells was stained with 100 nM Calcein-AM dye diluted in HBSS for 5 minutes on ice at a cell concentration of 1 million cells per milliliter. To potentiate intracellular esterases activity, cells were incubated for an additional 20 minutes at 37°C. Centrifuging at 240 g for 7 minutes, cells were washed twice with PBS and finally resuspended in media. To ensure cell contact, 75,000 of each Tag-it Violet and Calcein labeled cells were combined in the wells of a 96 well flat bottom plate in media containing either 5 mM 1-Heptanol or vehicle control and incubated for 2 hours at 37°C. Calcein transfer to Tag-it Violet+ cells was measured by flow cytometry by comparing to unstained controls.

### RGD peptide adherence assay

B16 cells were suspended in cold B16 media with or without 1 mg/ml RGD peptides. Two hundred and fifty thousand cells were then plated per chamber of an 8-chamber collagen I coated microscope slide (Corning, #354630). Cells were incubated for 45 minutes at 37°C and 5% CO_2_. After incubation, the media was removed and replaced with PBS, and cells were imaged. The PBS was then removed and combined with initial media as the non-adherent fraction, and 200 μl warmed Trypsin-EDTA (0.25%) was added to each well and incubated for 5 minutes. Two hundred microliters of media were then added to each chamber and pipetted up and down to ensure removal of all adherent cells. This was considered the adherent fraction.

### Statistics

Data were analyzed by unpaired two-tailed Student’s *t* test for parametric data, and one way analysis of variance (ANOVA) followed by Tukey’s post hoc test for multiple comparisons. Data normality determined using Shapiro-Wilks test. GraphPad Prism software was used for the analyses (GraphPad Software, La Jolla, CA).

## ACKNOWLEDGMENTS

We thank the BioImaging Shared Resources, Flow Cytometry and Laboratory Animal Facilities at Sidney Kimmel Cancer Center for their expert assistance in performing the presented experiments. We are very grateful to Dr. Hans Acha-Orbea for sharing with us the MutuDC cell lines. We thank the Sigal lab at Thomas Jefferson University for sharing reagents and mice with us. We thank Dr. Zoltán Fehérvári at Nature Publishing for critically reading the manuscript. Some of the figures were generated using BioRender.

## Author contributions

C.H. and B.Z.I. designed experiments. C.H. performed most of the experiments, analyzed the data and generated the figures. Q.S. made the original observation that conditional deletion of *Gja1* in LCs does not lead to deficiency. A.B. confirmed the experiments performed by Q.S. E.J.M. performed the cross-presentation experiments. Z.Q. did the allogeneic ICM experiments. D.M.B. performed the optimization of human DC differentiation from CD14+ blood monocytes. E.G. performed bioinformatic studies. C.H. and B.Z.I. wrote and edited the manuscript. All authors participated in discussions of experimental results and edited the manuscript.

## Data and materials availability

All data needed to evaluate the conclusions in the paper are present in the paper or the Supplementary Materials.

---

**Supplemental Fig. 1.**
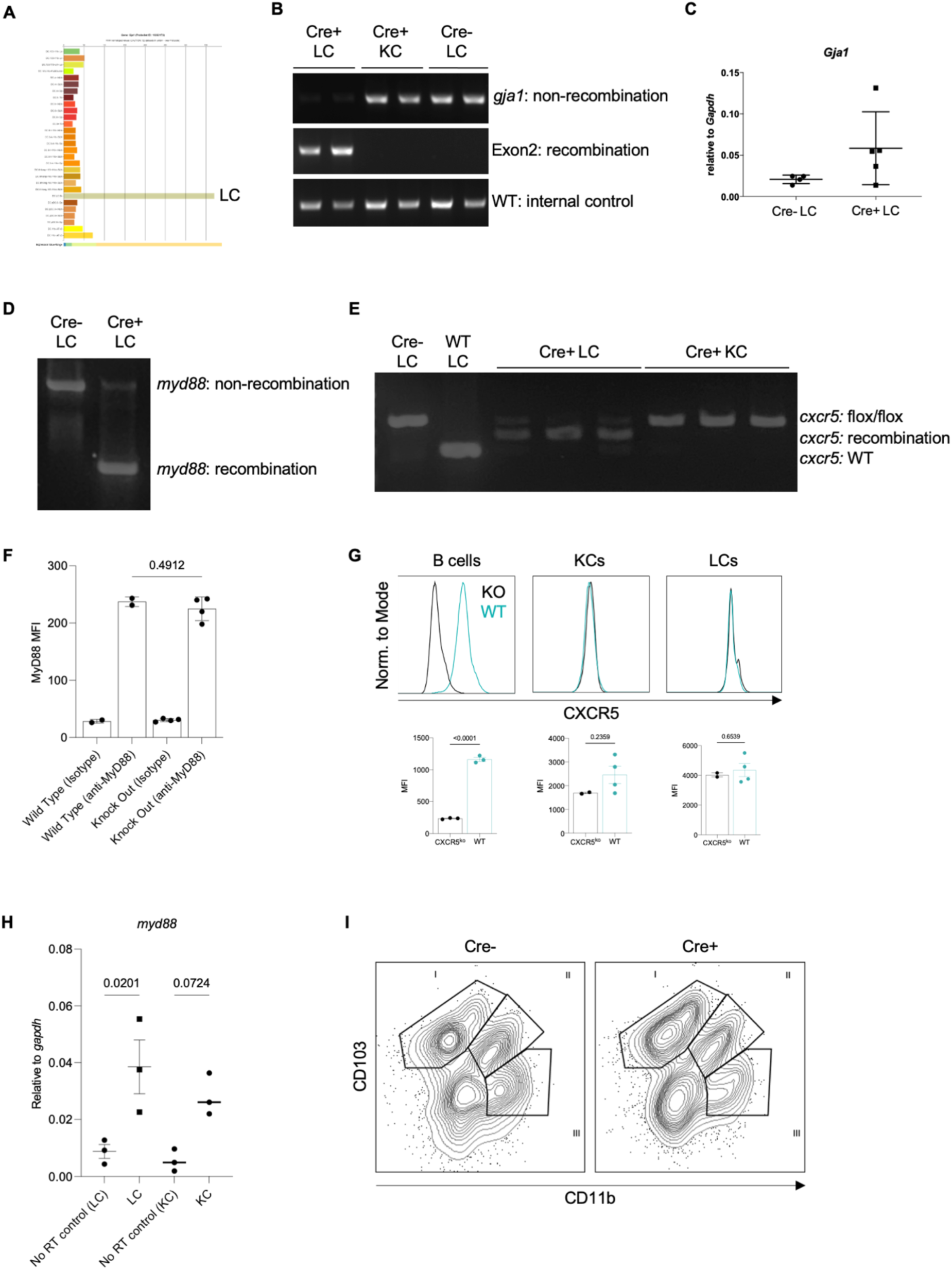
(**A**) Results of a *gja1* (Cx43) query into the ImmGen microarray database showing levels of *gja1* RNA transcripts in dendritic cell subsets. (**B**) Successful genetic recombination of *gja1* (Cx43) locus in Cre+ LCs, but not KCs or Cre- LCs. Cells were sorted from the epidermis of Cre positive and negative mice. (**C**) *gja1* mRNA levels in LCs sorted from Cre negative and positive mice. Each dot represents a separate mouse. (**D**) Successful genetic recombination of *myd88* locus in Cre+ LCs, but not Cre- LCs. Cells were sorted from the epidermis of Cre positive and negative mice. (**E**) Successful genetic recombination of *cxcr5* locus in LCs sorted from Cre+ mice. (**F**) Intracellular MyD88 flow staining of whole blood from wild type or MyD88 global knock out mice compared to isotype control. Dots represent individual mice. (**G**) CXCR5 staining of B cells, KCs, or LCs from CXCR5 global knock out mice or wild type mice and summary graphs. Dots represent individual mice. (**H**) *myd88* mRNA levels relative to housekeeping gene *gapdh* in LCs and KCs sorted from wild type mice. Dots represent individual mice (I) Example flow plots showing CD11b+CD103+ mesenteric lymph node DCs (II).

**Supplemental Fig. 2.**
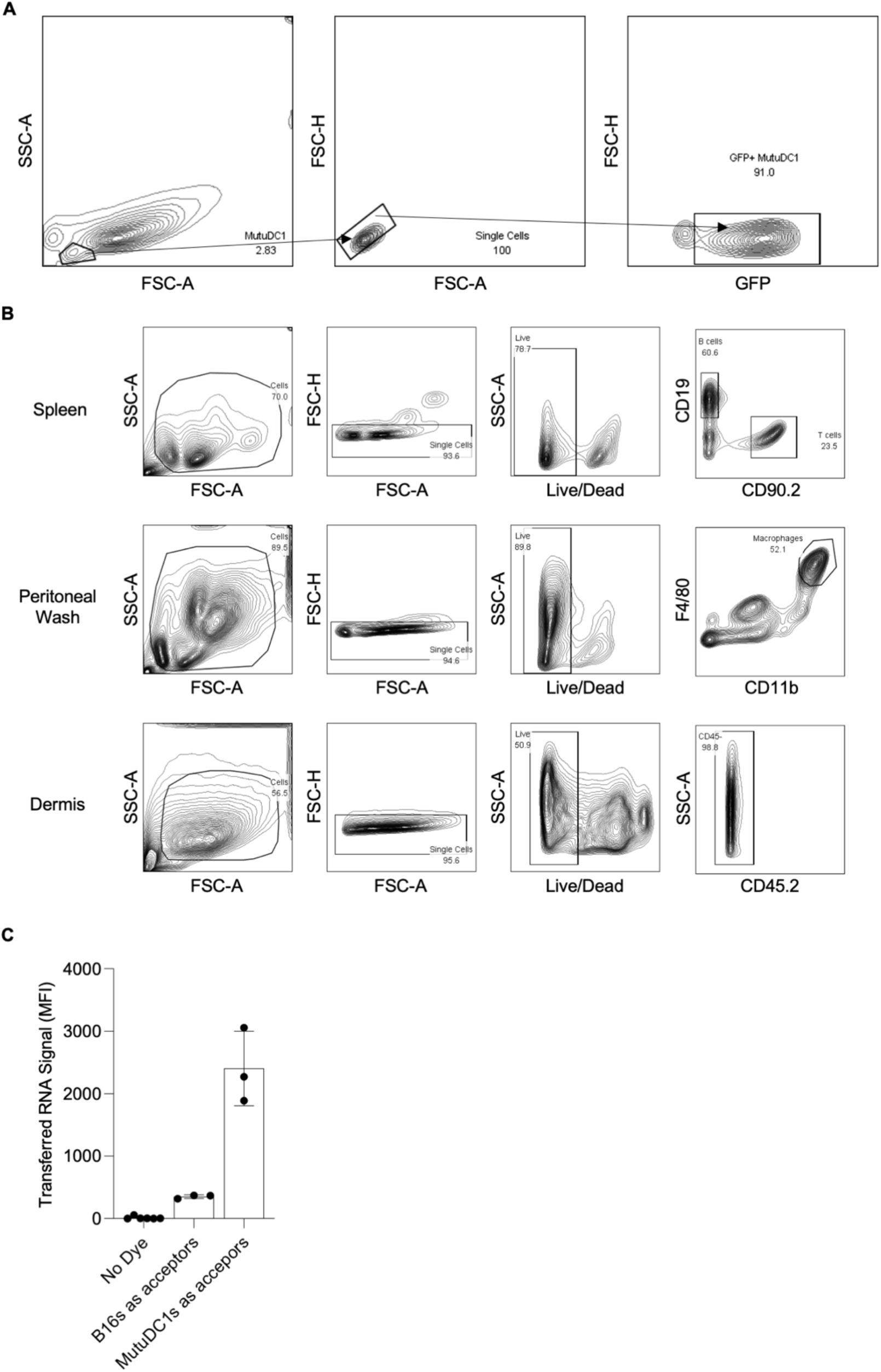
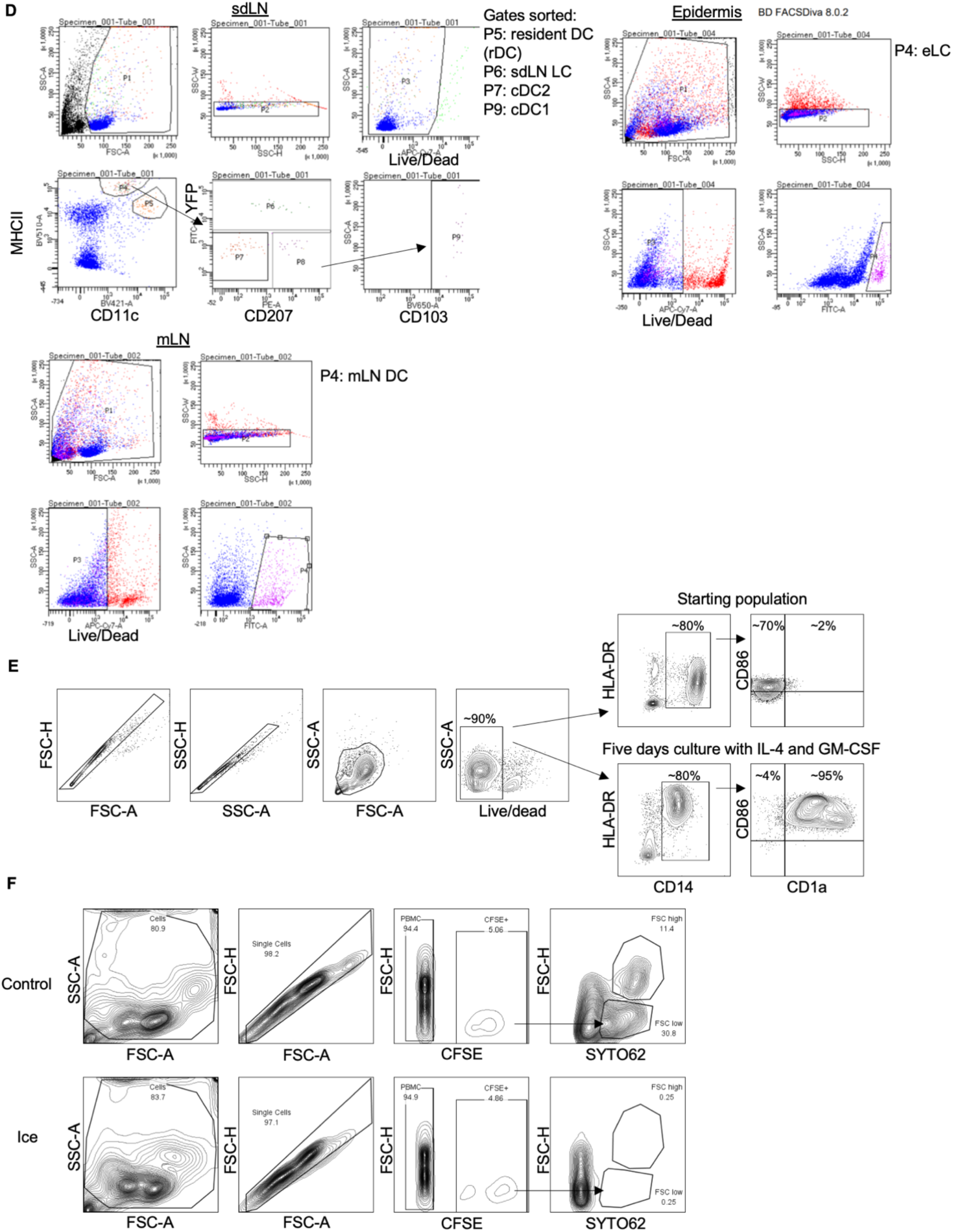
(**A**) Gating strategy for distinguishing GFP+ or CFSE positive cells after co-culture. (**B**) Gating strategy for sorting splenic T and B cells, peritoneal macrophages, and dermal CD45- cells. (**C**) Comparison of the transferred RNA signal contained within B16s or DCs. Either B16s or DCs were RNA labeled then co-cultured with the other cell type for 45 minutes at a 1:1 ratio. Dots represent individual replicates. (**D**) Gating strategy for sorting epidermal Langerhans cells, skin draining lymph node migratory Langerhans cells, sdLN cDC1, cDC2, and resident DCs, and mesenteric lymph node migratory DC from Cre+ hLangCre-YFP^f/f^ mice. (**E**) Differentiation of human DCs from blood monocytes. Gating strategy. One representative experiment out of two is shown. (**F**) Gating strategy for identifying CFSE labeled DCs in DC/PBMC co-cultures. Summary plots calculated with FSC-H high populations. One representative experiment out of two is shown.

**Supplemental Fig. 3.**
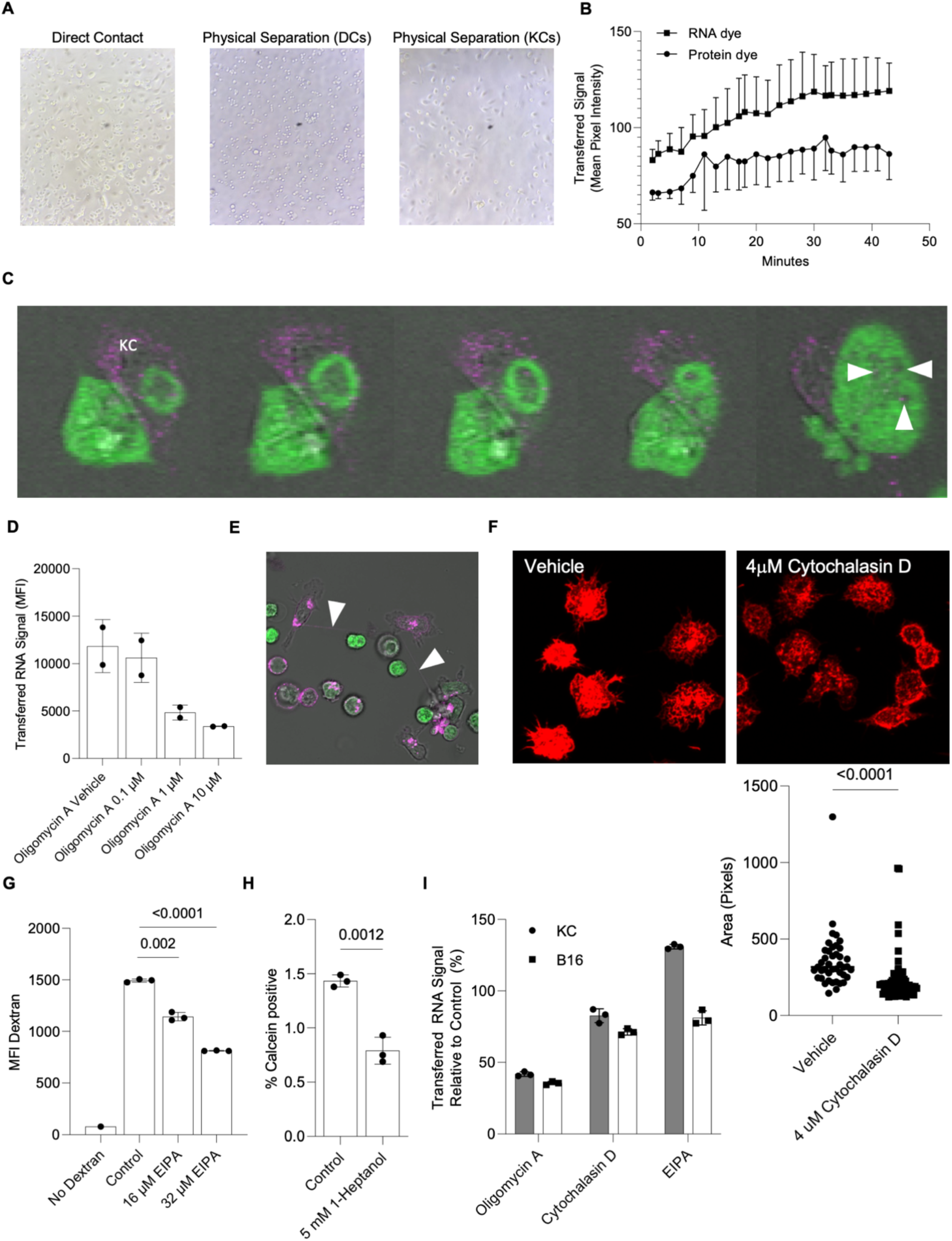
(**A**) Representative images of: MutuDC1/keratinocyte direct co-culture, MutuDC1 cells on the bottom of the well, or keratinocytes adherent to the 8 mm coverslip while suspended overtop of MutuDC1s. (**B**) Mean pixel intensity of transferred RNA or CTV signals contained within areas identified to be dendritic cells over a 45 minute live cell time lapse. Each point represents the average of 4 fields of view of RNA or CTV signal contained within DC areas. (**C**) Confocal single Z-plane slice of MutuDC1 cell (green) interacting with SYTO62 stained keratinocyte (purple). White arrowheads point to RNA containing vesicles in DCs. (**D**) Transferred RNA signal contained within MutuDC1 cells after a 45 minute co-culture with RNA labeled KCs containing indicated concentrations of Oligomycin A. (**E**) Example of TnTs (white arrowheads) forming between DCs (purple) in DC/KC co-cultures. (**F**) Rhodamine-Phalloidin staining of MutuDC1s in the presence of absence of 4 µM Cytochalasin D. Cell area measurements of individual cells in the presence or absence of 4 µM Cytochalasin D. Dots represent individual cells from a single experiment. (**G**) MutuDC1 TRITC mean fluorescence intensity after a 30 minute incubation in media containing 10 kD TRITC Dextran, 16 µM EIPA, 32 µM EIPA, or vehicle control. (**H**) Percent Calcein positive acceptor KCs (CTV labeled) in the presence or absence of 5 mM 1-heptanol. Dots represent individual replicates. (**I**) Transferred RNA signal contained within MutuDC1 cells after a 45 minute co-culture with either RNA labeled KCs or B16s in media containing 1 µM Oligomycin A, 8 µM Cytochalasin D, or 32 µM EIPA. Dots represent individual replicates.

**Supplemental Fig. 4.**
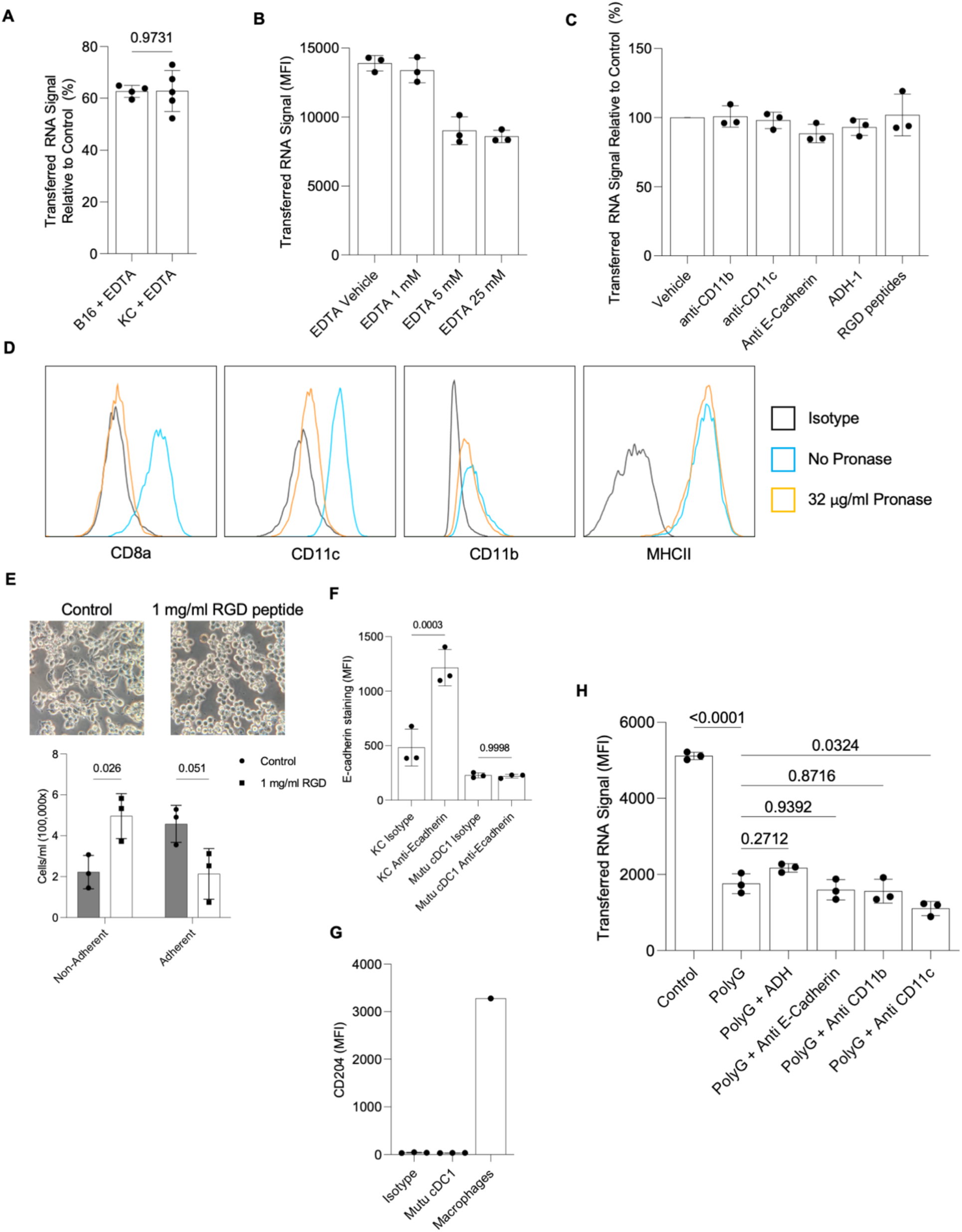
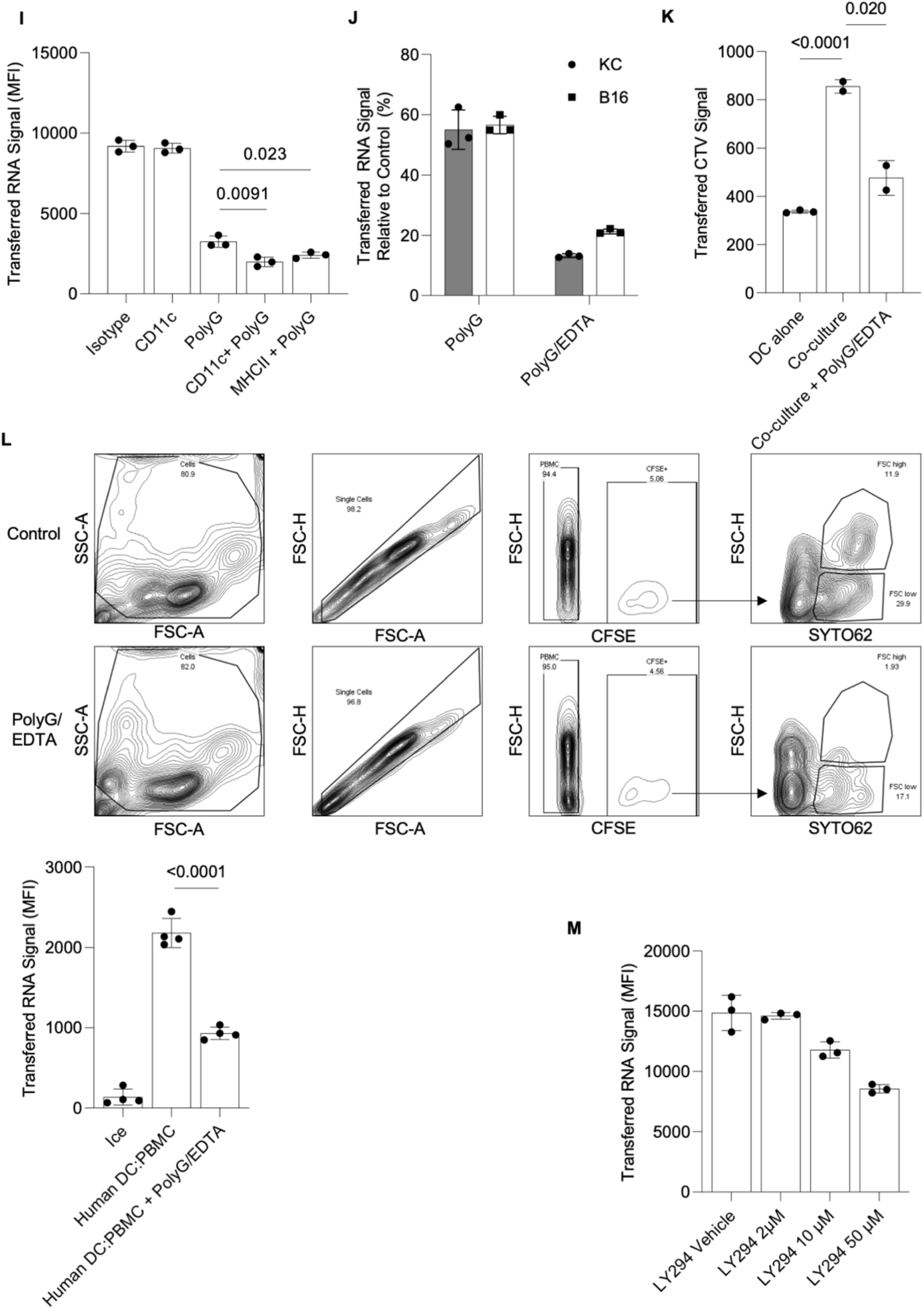
(**A**) Percent inhibition compared to control of RNA signal contained within MutuDC1s after a 45 minute co-culture with either RNA labeled keratinocytes or B16 cells in the presence of 5 mM EDTA. (**B**) RNA signal contained within MutuDC1 cells after a 45 minute co-culture with RNA labeled KC containing indicated concentrations of EDTA. (**C**) Percent inhibition compared to control of RNA signal contained within MutuDC1s after a 45 minute co-culture with B16s in the presence of 2.5 µg/ml anti-CD11b blocking antibody, 2.5 µg/ml anti-CD11c blocking antibody, 5 µg/ml anti-E-cadherin blocking antibody, 350 nM ADH-1, or 1mg/ml RGD peptide. (**D**) Staining of MutuDC1 cells or MutuDC1 cells treated with 32 µg/ml Pronase and antibodies against the indicated proteins at a concentration of 2.5 µg/ml. (**E**) Number of adherent and non-adherent B16 cells after a 45 minute incubation on collagen coated microscopy slides in the presence or absence of 1 mg/ml RGD peptide and images of cell morphology just prior to removal of supernatant. (**F**) Binding of a blocking anti-E-cadherin antibody to either keratinocytes or MutuDC1s. Binding was detected using goat anti-rat IgG DyLight 550 conjugate antibody. (**G**) MutuDC1 cells or peritoneal macrophages stained with 1 µg/ml anti-CD204 antibody. Dots represent replicates taken from a single mouse. (**H**) RNA signal contained within MutuDC1 cells after a 45 minute co-culture with RNA labeled B16 cells in the presence of indicated inhibitors or antibodies plus 500 µg/ml PolyG. (**I**) RNA signal contained within MutuDC1 cells after a 45 minute co-culture with RNA labeled B16s in the presence of anti-CD11c or anti-MHC-II antibody + 500 µg/ml PolyG. (**J**) Transferred RNA signal relative to control contained within MutuDC1 cells after co-culture with either RNA labeled B16s or keratinocytes in the presence of 500 µg/ml PolyG or 500 µg/ml PolyG + 5 mM EDTA. (**K**) CTV signal measured within MutuDC1 cells after 4 hour co-culture with CTV labeled B16 cells in the presence or absence of 500 µg/ml PolyG + 5 mM EDTA. (**L**) Gating strategy for identifying CFSE labeled DCs in DC/PBMC co-cultures. Summary plots calculated with FSC-H high CFSE+ populations. One representative experiment out of two is shown. (**M**) RNA signal contained within MutuDC1 cells after a 45 minute co-culture with RNA labeled B16s containing indicated concentrations of the PI3K inhibitor LY294002.

**Supplemental Fig. 5.**
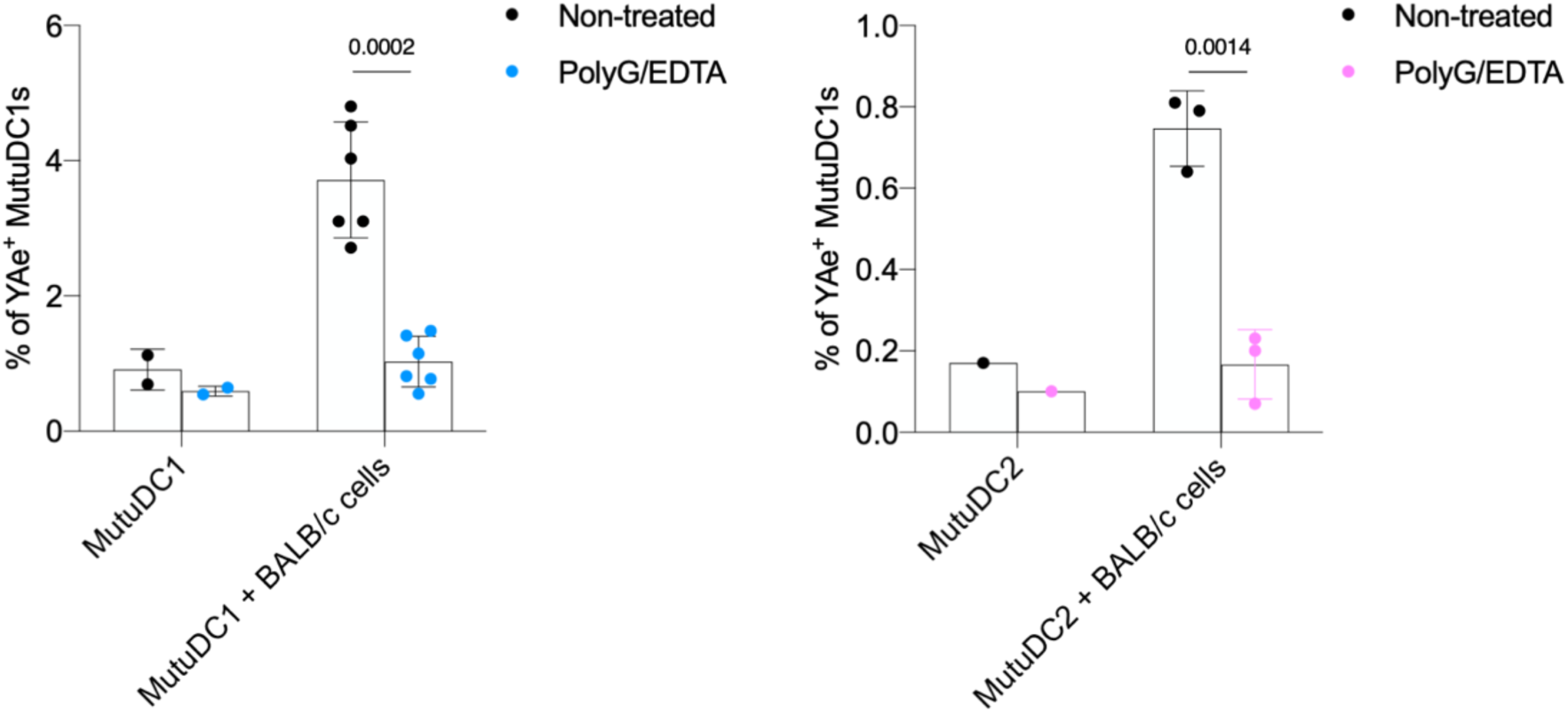
As in Fig. 6, MutuDC1 and MutuDC2 were co-cultured for 3 hours with BALB/c B cells in the absence or presence of PolyG/EDTA and then the Ea presentation of MHC-II determined by flow cytometry. Summary graphs for MutuDC1 (left) and MutuDC2 (right). Data from two independent experiments with 3 technical replicates were pooled, except for MutuDC2 which are from one experiment.

